# Ultra-fast joint-genotyping with SparkGOR

**DOI:** 10.1101/2022.10.25.513331

**Authors:** Hákon Guðbjartsson, Hjalti Þór Ísleifsson, Bergur Ragnarsson, Raony Guimaraes, Haiguo Wu, Hildur Ólafsdóttir, Sigmar K. Stefánsson

## Abstract

**Motivation:** Our aim was to simplify and speedup joint-genotyping, from sequence based variation data of individual samples, while maintaining as high sensitivity and specificity as possible.

**Results:** We have leveraged versatile GOR data structures to store biallelic representations of variants and sequence read coverage in a very efficient way, allowing for very fast joint-genotyping that is an order of magnitude faster than any joint-genotyping method published to date. Furthermore, it can be easily extended and executed much faster in an incremental fashion. Concordance analysis based on the Genome In A Bottle (GIAB) samples shows favorable results when compared with the de-facto standard approach, using gVCF files and GATK joint-calling. Additionally, we have developed variant quality classification using XGBoost and variant training sets derived from the GIAB samples. The entire business logic is implemented efficiently and concisely in SparkGOR.

**Availability:** SparkGOR is open-source and freely available at https://github.com/gorpipe.

## 1 INTRODUCTION

Genetic association studies that test whether a given sequence variant has involvement in controlling a given phenotype rely on ever increasing cohort sizes, in order to increase the power to detect disease association. These studies now often utilize variations derived from sequence read data in order to detect more rare variants and other features not readily available from array based data [11]. An important part in this process is to perform so-called joint-genotyping; to organize the genotypes into data structures that are efficient for large-scale cohort analysis, to properly account for lack of data due to low sequence read coverage, and to leverage population statistics to estimate, filter and improve quality.

Many of the early short read sequence variation callers use a position based model following sequence read alignment [15] where a Bayesian statistical model based on pileup of sequence bases quality and genotype priors is used, e.g. SAMtools, SOAP, and GATK’s Unified Genotyper [19][16][17]. These callers are highly effective for calling SNPs but less effective for InDels due to their reliance on independent read alignments to a reference sequence. To address this, algorithms using haplotype realignment and local assembly have been developed, such as GATK’s HaplotypeCaller, FreeBayes, Isaac, Platypus [5][6][23][24], as well as algorithms that use de Bruijn-like graphs for de novo assembly [12][20]. Although these algorithms call InDels with greater accuracy, they do not scale well, due in part to exponential increase in complexity with the number of samples, making joint-calling challenging for large cohorts, but joint-calling can be used to improve quality by providing cohort based distributions for variant attributes and support variant quality re-calibration (VQSR).

In order to enable joint-calling of larger cohorts, the reference confidence model (RCM) has been developed [22], where in addition to likelihoods for all alleles explicitly observed in sample reads, the model generates the likelihood over the set of unobserved non-reference alleles and stores them in gVCF format for each sample [23]. The primary purpose of this is to allow downstream joint-analysis of multiple gVCF files to distinguish homozygous reference from the void of any read data at all. Additionally, this reduces the computational burden associated with the so-called *N+1 problem* for incremental joint-genotyping, as cohorts increase in size.

For very large cohort sizes, with tens of thousands of samples, practical performance issues, related to the necessary parallelism, indexing, and random access to the relevant genome shards of each sample, start to play an important role [11][14]. To address this, systems such as GLnexus [18] have been developed where the gVCF blocks from individual samples are stored in a RocksDB database. The GLnexus system has also been optimized [29] for single sample gVCF files generated with the DeepVariant caller [21], but they have a smaller storage footprint because of efficient quantization of the reference records.

Deep-learning methods such as DeepVariant have also been augmented with *new channels* to encode allele frequencies [2] and sequence reads from parents [13] and shown to improve precision and recall in single sample calls.

Pangenomes have been proposed to remove bias toward the reference genome. They incorporate prior information about variation, allowing read aligners to distinguish better between sequencing errors in reads and true sequence variation. Unlike de novo assembly algorithms, pangenomes represent sequence variation with respect to the reference genome, enabling direct access to its annotated biological features. PanGenie [7] leverages a haplotype-resolved pangenome reference together with k-mer counts from short-read sequencing and allows read mapping and genotype calling to be performed efficiently in a single step, however, currently it cannot be used to genotype very rare variants that are present only in the sample, but in none of the other reference haplotypes. GraphTyper [8][9] relies on global alignment to the linear reference sequence to assign reads to regions. It then locally realigns sequence reads from a region to a pangenome graph, uses the resulting alignments to efficiently update the graph, and concomitantly genotypes sequence variants. This method has been shown to be better suited for population-scale genotyping than other widely used pipelines and provide improved sensitivity and precision [11].

Here we present a joint-genotyping method implemented efficiently in SparkGOR [27]. Our pipeline accepts single sample gVCF-like input and generates pVCF-like output. By storing variants and sequence read coverage information in separate *bucketized* GORZ files [10], we obtain up to 20 fold storage footprint reduction, as compared with conventional gVCF files, structures that are easy to update and lend themselves well to parallel processing. Also, by converting multi-allelic locus based variant calls to bi-allelic variants, we simplify and speed up our joint-genotyping computation dramatically while maintaining excellent quality and concordance with GIAB samples [30]. Finally, we aggregate variant and sequence coverage attributes and train a supervised XGBoost [3] variant classifier, using training data derived from concordance with GIAB samples. We show how this efficient QC classifier, that segregates good and bad variants, is consistent with GATK’s best practice VQSR quality filtering approach. Finally, we outline how our approach can be used for very efficient incremental joint-genotyping, something that can be of importance in a setting where large databases are frequently updated with new samples.

## 2 METHODS

### 2.1 Evaluation data

To test our joint-calling pipeline, we used about 37 thousand samples that had already been sequenced with Illumina-Novaseq technology. These samples are mostly of Caucasian ethnicity and their target sequence depth was 30 reads. Additionally, we downloaded 7 GIAB samples [30] and their BAM files were sub sampled to reach the same mean depth. Also, we sequenced one of the GIAB samples, NA12878, which represents the same individual as the data downloaded for sample HG001. The Sentieon implementation of GATK [25],[26] was used to generate gVCF files from the BAM sequence read files. These gVCF files are therefore the starting point for our GOR based processing pipeline. For joint-calling, we used both pipelines to processed all the samples, including the GIAB samples, for chr22 only. Because of the cost, we only processed the entire genome with our GOR pipeline, but for comparison, we had an earlier Sentieon run processed genome wide, with all the samples except the 7 GIAB samples.

### 2.2 Converting gVCF to GOR files

In our supplementary material, query Example 1 shows the logic used to split up a gVCF file into biallelic variants and sequence read coverage segments. In particular, the create step for `#biallelevars#` shows the commands used to generate the biallelic variants. Noteworthy is the use of the `PIVOT` command to move the attribute-value based content in the INFO and FORMAT/DATA columns into separate columns. Then there are the commands `CALC ngt`, `SPLIT ngt`, `CALC thePL`, and `REPLACE pl`, that are used to convert the multi-allelic locus representation into biallelic format, where each row corresponds to a single allele.

The other noteworthy create step is for `#segcov#` which picks up the depth and segment size, rounds the depth as shown in `CALC rd`, such that little rounding is used where the depth is low, and then combines adjacent segments with equal depth, using the `SEGSPAN` command. This makes the downstream data processing faster because the segments are larger and fewer; reducing the file sizes as demonstrated inTable 1 shows.

**Table 1.** File sizes for two example gVCFs and corresponding GORZ files

| Filetype | PN1 |  |  | PN2 |  |  |
| --- | --- | --- | --- | --- | --- | --- |
|  | Filesize in bytes | % | Rows | Filesize in bytes | % | Rows |
| gvcf.gorz | 8,382,422,788 | 100.0 | 439,304,078 | 3,336,629,971 | 100.0 | 154,518,568 |
| biallelefile.gorz | 260,918,637 | 3.11 | 5,083,826 | 258,550,605 | 7.75 | 5,065,486 |
| segcovfile.gorz | 103,407,142 | 1.23 | 55,062,862 | 51,163,969 | 1.53 | 25,370,440 |
| lowcovfile.gorz | 7,265,032 | 0.09 | 3,369,573 | 5,903,302 | 0.18 | 2,709,705 |
| goodcov10.gorz | 1,795,958 | 0.02 | 651,303 | 1,419,577 | 0.04 | 603,447 |
| goodcov8.gorz | 1,503,959 | 0.02 | 609,043 | 1,221,080 | 0.04 | 575,143 |
| goodcov6.gorz | 1,236,350 | 0.01 | 571,480 | 1,042,059 | 0.03 | 559,519 |
| goodcov4.gorz | 985,993 | 0.01 | 548,203 | 862,595 | 0.03 | 542,570 |

The gVCF2GOR queries in Ex. 1 take about 10-15 minutes to run for a single gVCF file when executed in parallel. The overall computation cost is about 1.5 core-hour and while the script could be optimized by writing dedicated GOR commands to handle multiple pipe steps, as compared to using only standard general purpose GOR commands, the overhead of running the script is still only four to five times that of reading and de-compressing the gVCF file. Since this is a one-time operation per gVCF file and much less expensive than the variant calling step for generating the gVCF file in the first place, we have not spent time on writing dedicated commands for this purpose. For comparison, running our Sentieon pipeline on a 36 core instance with 72 GB of memory, it typically uses 2.5 hours for BWA alignment, 12min for marking duplicates, 15min for realignment, 11min for BQSR, and 46min for the GATK-like haplotype variant calling step.

In Table 1, we see two examples of gVCF files and their corresponding GORZ derivatives as specified in query Ex. 1. The gVCF files are significantly different in size because of their difference in storing `<NON_REF>` records. We see that the GORZ files are in total only 1/20th to 1/10th of the original gVCF files. The biallelic variants take most of the storage because the coverage segments are highly compressed due to the rounding of the coverage depth, as mentioned before. As their name indicates, the “goodcov” files store just sparse binary segments, the presence of the segment indicating that the coverage exceeds a certain coverage threshold and their absence that the coverage is lower. Similarly, both the “segcovfile” and the “lowcovfile” are storing information on approximate coverage, the latter capping the depth on 15 reads; both representing zero depth in a sparse manner with the absence of segments.

In the joint-calling steps discussed later, we don’t need the full coverage information, but rather, we need to know it where it is relatively bad, in order to distinguish lack of data from homozygous reference. Therefore, in the downstream joint-calling step, we only need `biallelefile.gorz` and `lowcovfile.gorz`, making the data input less than 1/10th of gVCF input.

### 2.3 Bucketizing GOR tables

When each gVCF sample has been processed with the gVCF2GOR queries presented above, we insert them into GOR tables [10] which we typically refer to as GOR dictionaries or GORD files. The dictionaries are GOR relations that are partitioned and the GORD files are essentially metadata files pointing to the partitions that are selected using the `-f` or `-ff` filtering options on the `GOR` command. Unique to GOR dictionaries is that they support two partition levels, or so called buckets, providing very rapid table insert and read access in selective queries. The bucket partitions then enable efficient access (in terms of file driver memory footprint and random file seeks) in less selective cohort type queries and the `GOR` command will try to optimize which partitions to use for mixed scenarios. Furthermore, the parallel macro command `PARTGOR` provides an easy way to optimize distributed parallel queries based on storage partitions and input selection criteria, allowing for parallelism, not just across the genome as with `PGOR`, but also across the data partitions.

Query Ex. 2. in the supplementary material shows an example of how easy it is to create the bucket partitions, given that all the original sample level partitions are present in a file such as `biallele.gord`. With dual partitions the data footprint is approximately duplicated, with the benefit of having rapid access to data from few selected samples. If the data is only to be used by a joint-calling process, and not by ad-hoc queries with different selectivity, then only the bucket partitions are needed, making the storage comparison with gVCF slightly better than shown in Table 1, because increased compression ratio often observed in buckets due to the homogeneity of the data.

By using a bucket size of 500 as shown in our example the number of files, that are initially randomly accessed and subsequently simultaneously kept open for streaming for each parallel worker, is reduced by the same amount. This naturally eliminates the issues related to Tabix indexes, as discussed in [11][14][8][18], solved by them with custom index strategies for multiple samples or by using database systems, such as RocksDB or LevelDB, that support batch writing, somewhat analogous to the bucket strategy in GOR. Importantly, although not shown here, the bucketization process for GOR dictionary tables can be made fully incremental, and as long as the joint-genotyping process uses a substantial fraction of the data in the buckets, there is no need for re-bucketization.

### 2.4 Joint calling

The actual code for the joint genotyping step is shown in query Ex. 3. The implementation shown here uses a distributive model that can scale to extremely large number of samples and lends it well towards and incremental approach. This approach is analogous to the one used by GLnexus approach [18], i.e. where a subset of the samples are called together. Hence, it requires two passes through the variants; one to find all possible variants present in the sample population and then call each sample subgroup at every given variant.

There are two create steps, `#part_allvars#` and `#allvars#`, that find how many rows there are for each variant. The first step executes in parallel across the genome for each bucket partition and the second one aggregates the counts across the partitions. We then generate `#jc_buckets#` to define multiple groups of 5000 samples that are called together with the `PRGTGEN` command. This command is the only one presented in this work and our example queries that is not a general purpose GOR command and specifically written to speed up joint genotyping. This command takes a left-input stream of variants and joins them with a right-input stream of low coverage segments (see `#lowcov_dict#`). Additionally, it gets as an input the relation `[#jc_buckets#]` to specify how the genotypes should be arranged horizontally in a `values` column in the output rows. This is equivalent to different columns in pVCF files except that different horizontal buckets are stored in different row partitions and therefore amenable to incremental update. Note that this relation is filtered to include only the samples being processed in each group, because the input is assumed sparse and the absence of coverage row for a sample is equivalent to zero coverage and similarly the variants are sparse, i.e. only present where called and included by the `-ff` option in the `GOR` source commands. Also notice that the `-split` option is used to call variants in genomic segments as specified by `[#par_segs#]`, i.e. 1M variants per worker job as defined with the `SEGHIST` command.

The `PRGTGEN` command has multiple options and can either use sequence read depth and allele call-ratio or PL probability triplets to estimate the quality of the single sample called variant, the latter being preferred since it does not assume a constant base quality across the read pileup. It does also have options to specify the genotype priors in the single sample calling step (`-fpab` and `-fpbb`), the error rate per read, *ϵ*, and the genotype probability threshold, to avoid an “unknown” call. Also, if there is a close call between heterozygote and homozygote call, but neither crosses the threshold, the option `-combgt` can be used to combine these probabilities to see if the one more likely should be pushed over the threshold to avoid an unknown call, e.g. something that may be useful for gene burden analysis of rare variants.

For samples where there is no variant record, the coverage estimate in `#lowcov_dict#` is used to estimate the PL triplet as ((1 − *ϵ*)^*d*^,(1*/*2)^*d*^,*ϵ*^*d*^) for the genotypes (hom-ref,het,hom-alt), where *d* is the read count or depth. Typically, the value for *ϵ* is small and therefore 1 − *ϵ* ≈ 1 and the probability for the homozygous reference will quickly dominate in the triplet as *d* is increased. Thus the joint calling is not very sensitive to the accuracy of the *ϵ* estimate. Likewise, we argue that it is NOT important to distinguish between a sequence read depth of 15 or higher, nor with greater resolution for lower depths than the definition in Ex. 1 specifies, i.e. a resolution of two reads.

While the `PRGTGEN` command can accept additional input with genotype frequency priors (not shown in our example) we typically don’t see a great need for that if sufficient number of samples are called together, e.g. in our example we do it in batches of 5000 samples. Our command, like so many other systems, uses an iterative EM algorithm [4]; to estimate the sample genotypes based on the probability likelihood triplets and the genotype priors, and then to update the priors based on the estimated genotype frequencies. Now, one can show that with N samples the chances of detecting a variant with AF of 1*/*(2*N*) is only 1*/e* ≈ 37% and for 1/10th of that frequency it is ≈ 10%, for 1/100th ≈ 1%, and so on. Thus, when the samples are only a finite subset of a population, we risk not observing variants and estimating the prior as zero. Therefore, in our EM estimate, we add a correction factor to the genotype frequencies to avoid that they fully converge to zero. For a Bayesian model, the priors don’t matter that much, if there is significant amount of data, because then *P* (*D* | *G*) converges to zero for the incorrect genotypes, *G*, as the number of data points/reads grows. Indeed, the priors will mostly impact the decision between unknown and homozygous reference, in the spectrum of common and moderately common variants where the coverage and quality is poor.

Because the Bayesian EM logic in the `PRGTGEN` command uses only bialleic variants, it fits well with a streaming architecture, uses little memory, and is computationally fast. Contrast this with models that take into account multiple allele combinations per individual, where the likelihood state space grows quadratically with the number of possible alleles per loci. Therefore, algorithms like GATK and GraphTyper provide a parameter to prune and collapse alleles together to limit the number of possible genotypes, e.g. in InDel repeat regions, an approximation that is related to the biallelic approximation used here.

### 2.5 Generating variant attributes for QC

Although a typical WGS sample harbors between 4 and 5M variants, the total number of observed variants in joint genotyping can easily be several hundred millions, for tens of thousands of samples (e.g. 300M for 30k samples); mostly rare variants. Figure 1 shows how the number of distinct variants we observed on chr2 as a function of number samples; a relationship that seems to be quite close to the square-root of the sample number^1^.

**Fig. 1:**
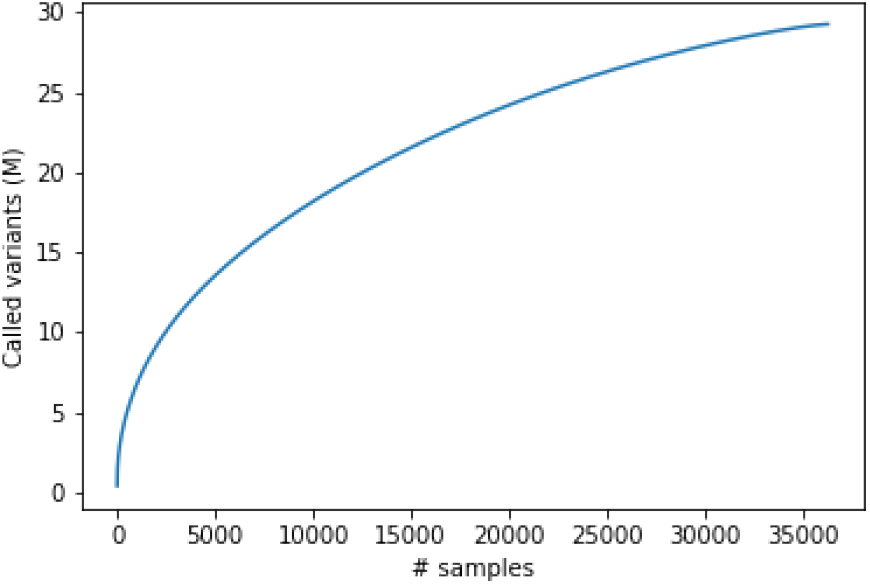
Observed variants on chr2 as a function of sample number.

In practice, many of these very rare variants may be due to sequencing errors or alignment imperfections. Therefore, it is very important to be able to classify the variants and filter out *bad* variants that may cause spurious associations in downstream GWAS analysis and distract from real signals.

From the genotypes, we aggregate standard metrics such as AF, HWE, yield etc. Example. 4 in the supplementary material shows example GOR queries that use the genotypes that we generated in Ex. 3 and stored in *horizontal bucket structures*. With high parallelism, this query runs quickly, because of the small data footprint of the genotypes and because it uses a counting command `CSVCC` that is designed and optimized for rows with horizontal values.

With the output from the previous query, we use the queries in Ex. 5 to evaluate *regional* attributes, based on adjacency to other variants and characteristics of the reference sequence. These are just few examples of easily calculated features and nowhere an exhaustive list of all meaningful possibilities.

Another set of relatively easily calculated features, that are unique to our approach, are shown in Ex. 6. There we calculate segments across the entire genome, with the total sequence read depth, in create step `#depth_segs#`, and segments with counts showing how many samples have coverage below a given threshold, e.g. `#lowCovQC4#`,`#lowCovQC6#`, …

Then we generate attributes based on the variant calls, that are similar to those generated by GATK, e.g. MD, QD, FS, SOR, BaseQRankSum, MQRankSum, and ReadPosRankSum. In Ex. 7, we show a query that reads through all the variants and aggregates metrics based on their annotations, such as those derived from the INFO field in the gVCF files.

At present, the query steps shown in Ex. 7, are the only ones that are written in a non-distributive manner and hence do not lend themselves to incremental update. This is because we use the median operator for some of the features, instead of average, to be compatible with the GATK specification. Also, we are just using the standard `GROUP` aggregation command and have not spent time generating an approximate distributive median calculation. Regardless, with all the features calculated, the query cost of create step `#varqc#` is still only 6 × higher than when only reading through the data. By creating a simple command that does multiple `CALC` steps and with some optimization for a specialized `GROUP` command, we could most likely cut the time in half.

Finally, inspired by [1] we also label regions and variants based on alignment quality from several samples. In Ex. 8, we show verbatim the query we used to calculate fbQ20, avg MapQ, and normalized average depth. This we did for 32 BAM files from our sample pool, but ensured that all the samples had a very close average read depth across the genome. Note that this BAM analysis is a one-time work that depends only on the sequencing method and the alignment steps and as long as they are unchanged, this does not have to be repeated. In Ex. 7, we can see the command `calc seqBad` where the number of bad labels are calculated, according to the table presented in [1].

### 2.6 Generating training data and GIAB concordance analysis

The thinned out GIAB BAM files were processed like our in-house samples and their gVCF files included in our sample processing pipeline, like the other 37k samples, and included both in our GOR joint-genotyping as well as our Sentieon implementation of the GATK-RCM pipeline [26]. For our concordance analysis, we ran the GATK pipeline only for chr22 while we ran our GOR *fast-freeze* pipeline described above for the complete genome. Then we downloaded the GIAB reference VCF files and the associated BED files that define the “trustworthy” regions. Example 9 shows how we process the files into corresponding GOR files. The logic for the VCFs is a simple version of Ex. 1 and the BED processing is only to ensure consistent order. We looked at the number of variants and the coverage of good regions with query as shown in Ex. 10 and found that all the BED files cover about 87% of the genome and the number of variants is close to 4M, thereof around 500k InDels.

In order to train a supervised classifier like XGBoost, we need positive and negative training samples. We use the concordance of GIAB samples for that purpose, i.e. comparison between the genotypes of these samples in the GOR joint-calling process and the genotypes observed in the reference VCF files, within the regions defined in the BED files. Notice that the BED files allow us to define false positive. The query logic in Ex. 11 shows the details of the analysis; `#GORvars_per_pn#` defines all observed variants, in either the reference files or the joint-called genotypes. These variants are joined with our observations and the truth and then we calculate pos-neg status as shown in `calc posneg`, after aggregating the truth status across the samples for each variant.

From `#posnegvars#`, we get 7,176,112 positive training variants and 684,020 negative variants; close to 10 × fewer. For testing our classifier, we leave out chr2 entirely from the training set and use it only for our AUC calculations, giving us 601,547 positive and 27,819 negative variants for scoring test. Also, for the training, we tried to balance the number of positive and negative training variants, as shown in the definition of `#training#`. This gave us 657,973 positive and 632,788 negative training samples.

The GIAB concordance analysis was performed for chr22. It compares GATK and GOR joint-calling genotypes, as well as the biallelic variants, with the GIAB reference VCF data and implemented in similar fashion as shown in Ex. 12. Notice that we count per `pn,callcopies,callcopiesx,depth,snp,good`, allowing us to calculate the sensitivity and precision in one genome sweep, for several scenarios of variant type; sequence read depth, and the variant classification that is implemented as discussed in next section.

Additionally, we calculate concordance for all the samples between genotypes produced with our GOR fast-freeze biallelic method and the GATK based approach. Query Ex. 16 in the supplementary material shows how we calculate the approximate mismatch rate for the variants and how many variants fall into each mismatch rate band. We define the mismatch rate in two ways; both based on all genotypes were neither samples has unknown call and for calls were either call in non-homozygous reference. This calculation is then done for both all variants, as well as only good variants, based on our variant classification as described below.

### 2.7 Training a variant classifier

We use a Spark implementation of XGBoost to train and predict variant classification. Ex. 13 shows the Python code that uses SparkGOR to define the dataframe `traindf`, using a parallel GORpipe query expression that generates a distributed RDD. As in regular Spark, all these manipulation of `traindf` are lazy until the model is evaluated in the very end. While this is not important for the training data, which in our case is only about 1.2M rows because of how few the GIAB samples are and the fact that we balance pos and neg variants, however, this is important for the prediction phase that may have to score many hundred millions of variants.

In addittion to scoring the variants with XGBoost based on the GIAB training data and the attributes shown in Ex. 13, we used the Sentieon release of the GATK VQSR Gaussian-mixture algorithm using the features MQ, QD, DP, MQRankSum, ReadPosRankSum, FS, SOR, InbreedingCoeff. This analysis we had performed earlier, using the entire genome with all the same 37k samples except for the 8 GIAB samples that we used for the concordance analysis, which as mentioned before was only evaluated for chr22.

We then compare the segregation power of these methods, using the pos/neg labels from the GIAB test data and calculate the AUC using the `roc_auc_score` function from the metrics module in the Scikit-learn Python package.

## 3 RESULTS

### 3.1 GIAB concordance comparison

The results from the concordance analysis are summarized in Table 2 for all the GIAB samples. We notice that the overall performance, as measured by F1, is best for our method implemented in GOR and similarly the overall sensitivity (TPR) and precision (PPV). Both of the joint-calling methods have better overall F1 score than the single-called (biallele), except for InDels, where GATK performs worse. These results are consistent with the fact that GATK joint-calling logic collapses variants in highly polymorphic sites and overall our results are consistent with GIAB concordance numbers reported from joint calling with UKBB data [11].

**Table 2.**
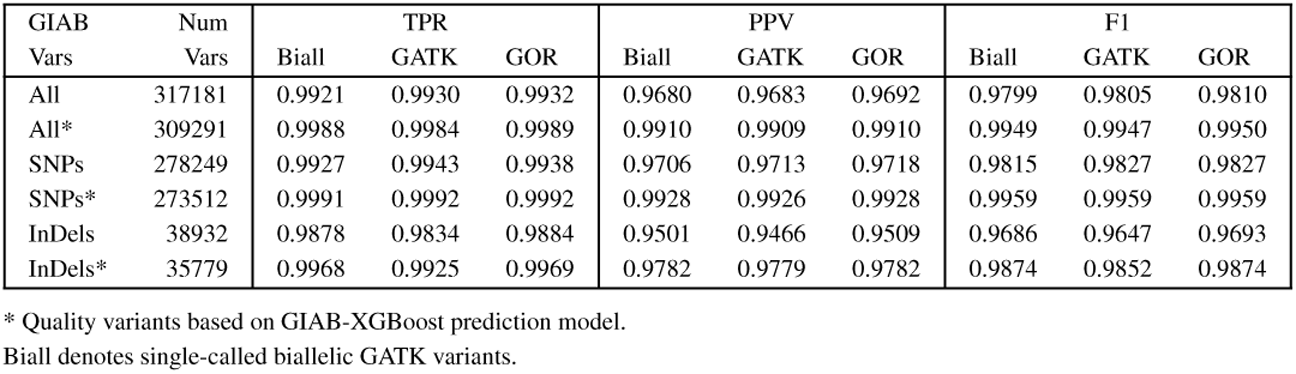
GIAB SNPs and InDels for chr22

| GIAB Vars | Num Vars | TPR |  |  | PPV |  |  | F1 |  |  |
| --- | --- | --- | --- | --- | --- | --- | --- | --- | --- | --- |
|  |  | Biall | GATK | GOR | Biall | GATK | GOR | Biall | GATK | GOR |
| All | 317181 | 0.9921 | 0.9930 | 0.9932 | 0.9680 | 0.9683 | 0.9692 | 0.9799 | 0.9805 | 0.9810 |
| All* | 309291 | 0.9988 | 0.9984 | 0.9989 | 0.9910 | 0.9909 | 0.9910 | 0.9949 | 0.9947 | 0.9950 |
| SNPs | 278249 | 0.9927 | 0.9943 | 0.9938 | 0.9706 | 0.9713 | 0.9718 | 0.9815 | 0.9827 | 0.9827 |
| SNPs* | 273512 | 0.9991 | 0.9992 | 0.9992 | 0.9928 | 0.9926 | 0.9928 | 0.9959 | 0.9959 | 0.9959 |
| InDels | 38932 | 0.9878 | 0.9834 | 0.9884 | 0.9501 | 0.9466 | 0.9509 | 0.9686 | 0.9647 | 0.9693 |
| InDels* | 35779 | 0.9968 | 0.9925 | 0.9969 | 0.9782 | 0.9779 | 0.9782 | 0.9874 | 0.9852 | 0.9874 |

\* Quality variants based on GIAB-XGBoost prediction model.
Biall denotes single-called biallelic GATK variants.

Table 2 shows similar numbers, now only based on variants predicted to be “good”, according our GIAB-XGBoost variant classifier. We notice a small increase in the numbers, however, the deviation from perfect concordance is approximately four times smaller. Table 3 shows similar analysis, but now only for variants in regions where the sequence read coverage of the samples was poor. Here we notice a great difference in the number, with or without the variant quality filtering. Also, we see that GATK has a slight advantage for the SNPs without filtering, while our GOR methods performs better on the InDels and when filtering is applied. Single-called gives always the worst F1, except for quality filtered InDels, where GATK has the lowest F1 number.

**Table 3.**
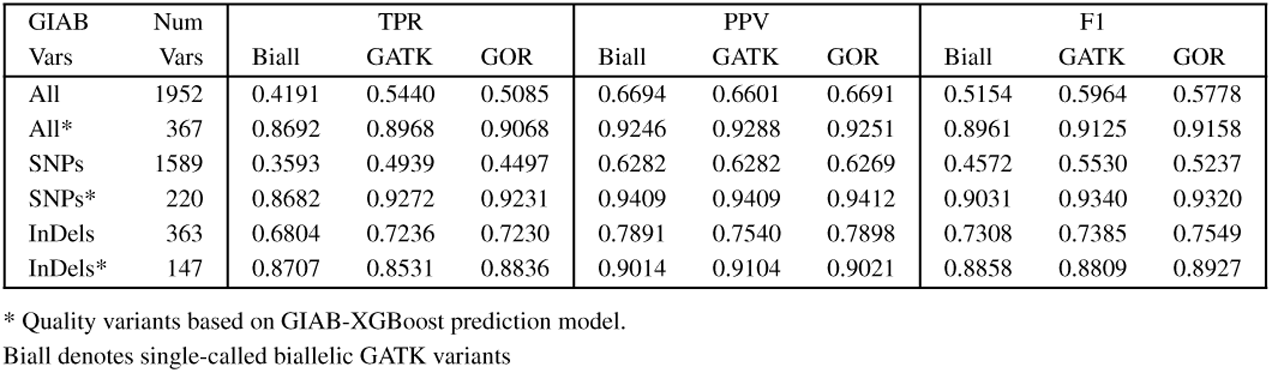
GIAB low cov SNPs and InDels for chr22, depth < 8

The supplementary material shows the same analysis for each sample as well as concordance numbers for our GOR method for the entire genome. There, we see that our joint-calling method gives consistently better F1 numbers than the single-called biallelic variants.

### 3.2 Variant classification

The variant classifications were carried out based on our XGBoost model trained on our GIAB training set, as described in section 2.6, and using a VQSLOD score derived using the GATK-VQSR process. Figures 3a-d) show the distributions for VQSLOD and the XGBoost score for a balanced set of variants on chr2; “pos” and “neg” defined based on the concordance of our GIAB samples as shown in query Ex. 11 in the supplemental material.

We notice a much more complex distributions for the VQSLOD than the XGBoost score and significant difference between the SNP and InDel distributions. For InDels, there is clearly limited segregation power in the VQSLOD score. The XGBoost score distributions are more similar although we see less certainty in the quality prediction for InDels.

The power of these distributions to segregate good and bad variants can be summarized using the area under the curve of receiver operating characteristic, ROC-AUC [28]. Table 4 shows the AUC value for these distributions based on all available test variants on chr2. We see that the XGBoost distribution has significantly higher segregation power, especially for InDels. For comparison, we also trained a second version XGBoost, using only the variant features used for the VQSLOD model. Interestingly, it compares quite well with the model using much larger number of features, although it does significantly worse for InDels. Overall, it is only 30% further away from the “perfect” classifier and shows that the features used for the GATK-VQSLOD model are highly informative.

**Table 4.** AUC for all test variants on chr2, defined using GIAB.

| Variants | Count | GATK-VQSLOD | GIAB-XGBoost | XGBoost v2 |
| --- | --- | --- | --- | --- |
|  |  | AUC | AUC | AUC |
| All | 629,366 | 0.919 | 0.981 | 0.975 |
| SNPs | 525,194 | 0.937 | 0.987 | 0.987 |
| InDels | 104,172 | 0.788 | 0.925 | 0.873 |

Typically, we have used a VQSLOD threshold of 2.41 to qualify good SNPs and a threshold of -1.77 for InDels^2^. For the XGBoost score we simply use a threshold of 0.5. Table 5 shows the correspondence between the variant classification using these two methods. Our GIAB-XGBoost based method eliminates 19% of the variants giving us 21M good variant on chr2 while with VQSLOD we eliminate 15%, keeping 22M variants as good.

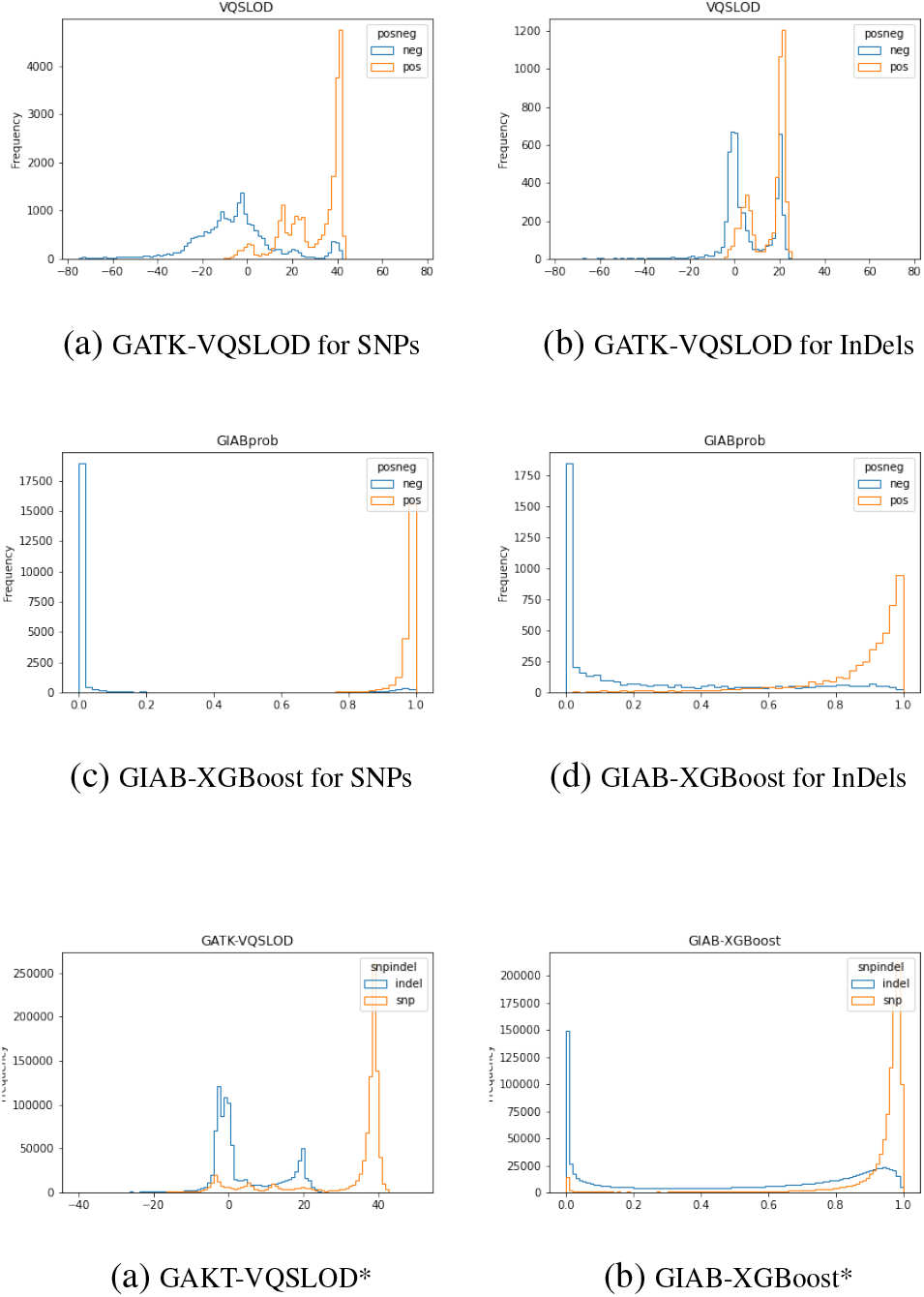
* Variants selected from chr2:50Mb-150Mb; 1M SNPs and 1M InDels.

**Table 5.** Quality labeling comparison for chr2

| GIAB | GATK | All |  | SNPs |  | InDels |  |
| --- | --- | --- | --- | --- | --- | --- | --- |
| XGBoost | VQSLOD | Variants | Fraction % | Variants | Fraction % | Variants | Fraction % |
| bad | bad | 2678794 | 10.27 | 2221934 | 9.61 | 456860 | 15.42 |
| bad | good | 2276259 | 8.73 | 1855395 | 8.03 | 420864 | 14.21 |
| good | bad | 1213982 | 4.65 | 473139 | 2.05 | 740843 | 25.01 |
| good | good | 19910484 | 76.35 | 18566887 | 80.32 | 1343597 | 45.36 |

### 3.3 Concordance between GOR and GATK

Tables 7 and 8 show results from the concordance analysis between GOR and GATK processes for all samples. Importantly, we do provide these numbers to reflect on the similarity between methods, but not to reflect on quality, unlike the numbers presented in section 3.1. The first table outlines the distribution of mismatch rate in all variants and the second show the analysis only for variants classified as “good” using our GIAB-XGBoost classifier. Both tables show two types of analysis. The left numbers based on dis-concordance rate where the denominator ignores genotypes where they are both homozygous reference^3^ and in the right column the mismatch rate calculation includes these genotypes^4^. We refer to these two different definitions of the dis-concordance rate as A1 and A2. The A2 analysis shows that less than 1% of the variants have genotype mismatch rate above 0.3% and for good variants the corresponding mismatch rate cutoff falls to 0.03%. When we define mismatch rate only from the non-ref calls, analysis A1, the mismatch rate rises. Still, less than 1% of the good variants have mismatch rate higher than 0.3%.

**Table 6.** Quality labeling comparison for chr2, with AF >= 0.001

| GIAB | GATK | All |  | SNPs |  | InDels |  |
| --- | --- | --- | --- | --- | --- | --- | --- |
| XGBoost | VQSLOD | Variants | Fraction % | Variants | Fraction % | Variants | Fraction % |
| bad | bad | 397669 | 17.53 | 348561 | 19.77 | 49108 | 9.70 |
| bad | good | 89168 | 3.93 | 83657 | 4.75 | 5511 | 1.09 |
| good | bad | 237714 | 10.48 | 25623 | 1.45 | 212091 | 41.91 |
| good | good | 1544237 | 68.06 | 1304831 | 74.03 | 239406 | 47.30 |

**Table 7.**
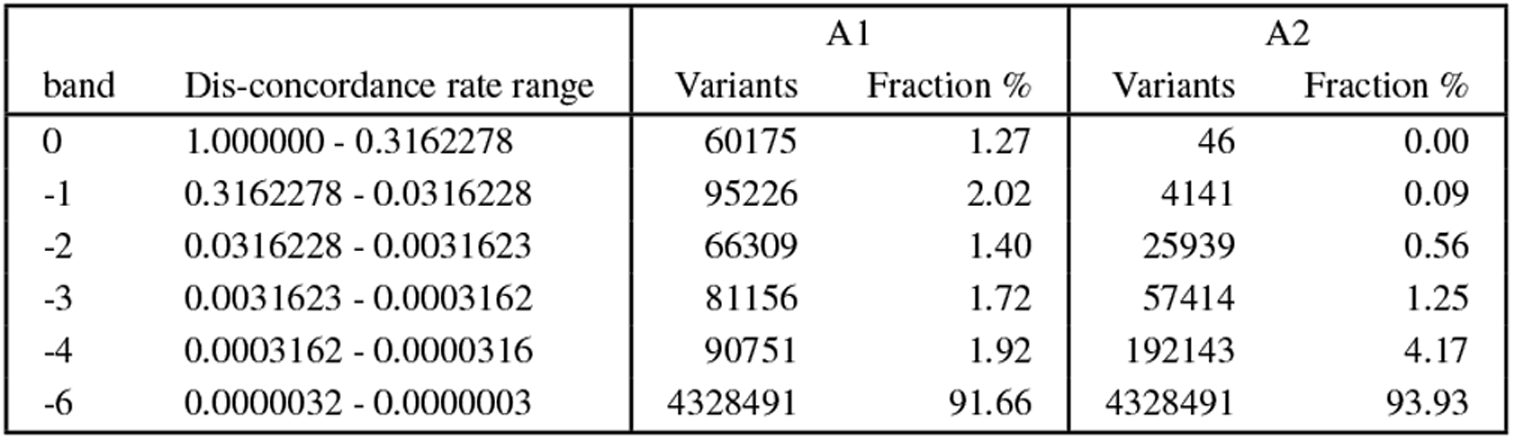
GATK-GOR genotype concordance for all variants on chr22

**Table 8.** GATK-GOR genotype concordance for “good” variants on chr22

| band | Dis-concordance rate range | A1 |  | A2 |  |
| --- | --- | --- | --- | --- | --- |
|  |  | Variants | Fraction % | Variants | Fraction % |
| 0 | 1.0000000 - 0.3162278 | 5405 | 0.16 | 1 | 0.00 |
| -1 | 0.3162278 - 0.0316228 | 4772 | 0.14 | 196 | 0.01 |
| -2 | 0.0316228 - 0.0031623 | 14189 | 0.42 | 2030 | 0.06 |
| -3 | 0.0031623 - 0.0003162 | 47493 | 1.42 | 8284 | 0.25 |
| -4 | 0.0003162 - 0.0000316 | 70184 | 2.10 | 88607 | 2.68 |
| -6 | 0.0000032 - 0.0000003 | 3201558 | 95.75 | 3201558 | 97.00 |

**Table 9.** Comparison of computation resources and cost estimates for 37k sample join-calling

| Method | Core-hours | Cost estimate |
| --- | --- | --- |
| GOR biallelic | 8.1k | \$370 |
| UKBB GATK4 (scaled estimate) | 1.1M | \$49k |
| Our inhouse Sentieon | 250k | \$11k |

### 3.4 Execution speed

As we show in the supplementary material, to generate the joint-calling genotypes, there are are several step involved in our processing of the single called gVCF output. The conversion from gVCF to GOR in Ex. 1 and the bucketization in Ex. 2 are one time processing steps that can be utilized for multiple subsequent joint-calling analysis. Therefore we do not consider them in the overall processing time, but for 37k samples it is about 55k core-hours for the complete genome.

In Ex. 3, the step to find all variants for the complete genome is about 450 core-hours. The joint-calling variant step itself, involving `PRGTGEN`, is only about 1500 core hours. However, the query in Ex. 7 that generates the variant attributes that are used for the variant classifier takes about 1000 core-hours. The queries in Ex. 4 and 5 are in total less than 250 core-hours and thus insignificant in the overall cost. The coverage query in Ex. 6 does however take 4900 core-hours, with the majority of the time, 4300 core-hours, spent on calculating the total depth across the genome, using `#segcov#`. The majority of the time is therefore not spent on the joint-calling step itself, but rather queries to gather metadata on the variants and their genotypes. This is a result of the fact that we are using general purpose GOR commands to aggregate these metrics and this logic is not folded into the `PRGTGEN` command itself^5^. For comparison, in query Ex. 3, we replaced the low coverage dictionary table with the segment coverage table. This resulted in 3.4× longer computation, i.e 4500 core-hours for the `PRGTGEN` step. This is because, as Table 1 shows, the segment coverage files are about 10× larger than the low coverage files and have over ten times as many rows.

The above numbers show that to generate joint-called genotypes for 37k samples and about 357M variants, the total computational cost with our biallelic GOR approach is 8100 core-hours^6^. Typical cloud computation cost for this is $370 and less than $200 by using spot instances.

For comparison, our cost for running the Sentieon-GATK pipeline for the same set is more than 30× larger! For alternative comparison, the numbers reported for calling 125k samples from UKBB using improved version of GATK4 [11] is 8.1M core-hours.

Scaling this number down by (35k samples * 357M vars)/(125k samples * 711M var) we get 0.14 * 8.1M core-hours, that is 1134k/13.8k = 82 times slower than our approach. Hence, we can infer that our joint-calling method is also significantly faster than the GLnexus joint-calling approach [29], which is reported to be 8 times faster than comparable GATK tools.

It is important to realize that in making the cost comparison based on cpu-hours, we are assuming it to be fixed across machine types and ignoring the memory requirements of the algorithms. Without going into detailed analysis of this aspect of the computation, we would like to mention that the GOR approach is very memory efficient, meaning that it does not need high memory/cpu ratio. While we can easily customize it, we typically use nodes with 36 cores and 64Gb memory for 28 GOR-workers^7^ and have never run into memory issues with our pipeline, in contrast with our experience executing our Sentieon-GATK pipeline. As we mentioned earlier, the variant aggregation step in Ex. 7 is the only step now that is not distributive in nature and hence the only step that may need larger memory as the number of samples grows. Alternativey, as we discussed before, we can approximate the median operator or replace is with the mean operator to solve this issue.

Here we have not listed the execution times for the XGBoost logic in Ex. 13, because it is small fraction of the overall computing time; usually executed in few minutes in Spark. Also, we have not included the time used for running the GATK-VQSR logic.

## 4 DISCUSSION

Here we have presented a very fast method for joint-genotyping large volume of samples and shown the entire code that is needed to implement and run it in parallel using SparkGOR. Our approach uses generic GOR files to store biallelic variants and sequence read coverage in separate files, therefore obtaining very high compression rates. We show that by simplifying the Bayesian joint-calling statistical model to deal only with biallelic variants and low-coverage approximations, we get very fast execution while preserving quality. By including the 7 GIAB reference samples in the joint-calling process, we have shown concordance, that exceeds what we observe for single-called samples and joint-calling with the GATK-RCM [22][26] method.

Additionally, we have presented a supervised XGBoost method for classifying variants based on multiple aggregate variant metrics and genome regional metrics. We then defined variant data sets with positive and negative variants, defined by the overall concordance of the GIAB samples. We compared this classifier with the GATK-VQSR approach, which is a semi-supervised clustering approach based on Gaussian mixture models. Our results show that our XGBoost approach yields better variant classification, based on the GIAB test data. While strikingly different approaches for classification, they do nevertheless result in similar numbers of good variants. Our classifier uses a rich set of features, many of which can be informative to humans when they explore variants or regions. However, here we have clearly not exhausted the number of possible features not done a thorough analysis of their importance. Neither have we done a rigorous exploration or automated search for optimal parameters to use in the joint-calling step itself, e.g. like presented in ref. [29], and is something that awaits future research.

At first, one might think that by feeding the joint-calling step only biallelic representations of the variants, that significant information is lost. However, when doing so, we are always picking the most likely single-called allele and only very rarely does some alternative genotype in the gVCF become more probable due to ensemble statistics. For instance, we have looked at double-heterozygous sites to see where our biallelic method could potentially give inconsistent genotypes (e.g. more than 2 allele copies per site). We have found that inconsistency occurs in less than one in million non-reference call genotypes and is fully absent in variants that are classified as good. Furthermore, although not done here, one should be able to provide the single-sample calling step approximate genotype frequencies, to make this a non-issue in the joint calling step. Providing population data has indeed been shown to improve quality in concordance comparison with GIAB [2].

By looking at the GIAB concordance numbers, we see that while our method gives the best F1 numbers, the different methods have quite comparable outcome for most of the variant classes. One can therefore argue that the primary purpose of a joint-genotyping step is therefore to distinguish between homozygous-reference and unknown, as well as to store the data in a format that is compact and efficient for regional based cohort analysis. Our output files are GORZ files, storing rows with genotypes in a value column, horizontally based on bucket structure. It has very similar data footprint as pVCF and PGEN files, but is more amenable for incremental update.

Indeed, as mentioned earlier, our method is easily extended to enable incremental joint-calling. As new samples are added to the pool, there will be new variants that have not been called before. These variants are easily identified by performing a negative join (`VARJOIN -n`) between a distinct list of all the variants in the new samples and the previous list of all distinct variants, e.g. `#allvars#` in Ex. 3. If we look at step `#joint_calls#`, for our previous samples, we can simply skip reading the biallelic variants, `#biallele_dict#`, because none of the new variants will be present there, and replace `#allvars#` with this new delta variant list. Thus, for all the old samples, we are only reading `#lowcov_dict#`, a table that is more than 25 times smaller than the biallelic variant input table, and performing joint-calling at very few sites compared with the total variant list. Using GOR dictionaries, we can also write the new genotypes called for the older samples into separate files, making incremental write trivial. This approach can be 100*×* more efficient than full genotyping, all depending on the ratio of new samples vs old samples. All the steps in our implementation, except for those in Ex. 7, are distributive in nature^8^ and can therefore be adjusted for incremental update. So far, we have not implemented the whole process to allow for incremental update since our full-update approach is very fast. However, one can envision scenarios where this is desirable, such as for research hospitals where data is being added on a regular basis. Our logic can also easily be used to provide reliable updates of AF, that properly takes into account lack of variant data because of low sequence read coverage.

The distributive nature of our approach provides it inherent scalability and option for incremental updates. We can also further increase the computational efficiency and speed of our approach by combining more business logic into commands, such as `PRGTGEN` which could for instance easily be modified to calculate the total read depth, given it is fed with the full coverage segments instead of the low coverage segments. By allowing us to drop steps involving `SEGPROJ`, this would lower cost of by 30% and by including more of the variant metadata logic, we could probably reduce the cost down to 50%. This will however reduce our agility in modifying our business logic and force deviation from the simple relational output model of our GOR commands, for moderate benefits. Some low level optimization of commands such as `SEGPROJ` might possibly achieve similar speed improvements, while allowing us to stay closer to our RISK-like Unix design philosophy, having a command set of “many small sharp tools”.

As a final remark, it should be evident that our method is quite generic and does not depend on the specific nature of the GATK gVCF files. Thus, our simple and efficient approach could be used to accelerate joint-genotyping by an order of magnitude, as compared to any other available method today, while maintaining high genotype quality.

## Supporting information

Supplemental query examples and tables.

## Footnotes

1 See Ex. 15 in supplementary notes for the query to generate this curve.

2 Corresponding to VQSRTrancheSNP99.90to100.00, i.e. 0.1% of the truth set is false positive.

3 See the filtering definition in `#temp1#` in Ex. 16, e.g. `gt1 != 3 and gt2 !=3 and not(gt1=0 and gt2=0)`

4 See definition of `#temp2#`, e.g. `gt1 != 3 and gt2 !=3.`

5 `PRGTGEN` was coded before we attended the variant scoring problem.

6 (450+1500+1000+250+4900) core-hours

7 Parallel queries in GOR are split up and executed on one or more GOR worker. A GOR worker executes one GORpipe query at the time but may use one or more threads to do so, depending on the commands in the query. A typical query uses about 1.2cores at the time.

8 Can be evaluated in batches of samples, e.g. with PARTGOR, and combined.

## Notes

### Competing Interest Statement

The authors have declared no competing interest.

### Summary of Updates

Minor edits; eliminating a deprecated URL and an incorrect citation.

