## Supplemental query examples and tables. for "Ultra-fast joint-genotyping with SparkGOR"

### Supplemental material for:
### "Ultra-fast joint-genotyping with SparkGOR"
### by Gudbjartsson, H. , et al.

#### 1 Scripts and query examples

**Example 1:** Converting gVCF file to GOR biallelic variants and coverage segments

```
def #filename# = s3://bucket/PN1.realigned.recalibrated.g.vcf.gz;
def #pn# = PN1;
def #writepath# = output_path/gVCF2GOR_files/#pn#;

def #lowcovfile# =  #writepath#/#pn#_lowcovfile.gorz;
def #segcovfile# =  #writepath#/#pn#_segcovfile.gorz;
def #biallelefile# = #writepath#/#pn#_biallelefile.gorz;
def #goodcov4# =  #writepath#/#pn#_goodcov4.gorz;
def #goodcov6# =  #writepath#/#pn#_goodcov6.gorz;
def #goodcov8# =  #writepath#/#pn#_goodcov8.gorz;
def #goodcov10# = #writepath#/#pn#_goodcov10.gorz;

def #filter_chrom# = where contains(chrom,'chr') and len(chrom)<6;

create #biallelevars# =  pgor <(gor #filename# | #filter_chrom#
| where alt != '<NON_REF>'
| rename #10 data
| replace info info+';GT='+vcfformattag(format,data,'GT')+';AD='+vcfformattag(format,data,'AD')
+';GQ='+vcfformattag(format,data,'GQ')+';PL='+vcfformattag(format,data,'PL')
+';SB='+vcfformattag(format,data,'SB')
| select 1-5,filter,info,qual
| split info -s ';' | colsplit info 2 x -s '=' | hide id,info
/* The below list may vary,  depending on variant caller.   Here, DB is willingly excluded */
| pivot x_1 -e 0 -v 'GT','PL','AD','GQ','SB','BaseQRankSum','ClippingRankSum','DP','ExcessHet','MLEAC','MLEAF',
'MQRankSum','RAW_MQ','ReadPosRankSum' -gc ref,alt,filter,qual
| rename (.*)_x_2 #{1}
| replace gt replace(gt,'|','/')
| where gt != '0/0'
| calc GL int(listnummin(listfilter(PL,'x!="0"')))
| calc ngt if(listfirst(gt,'/') != '0' and listfirst(gt,'/') != listlast(gt,'/'),'0/'+listfirst(gt,'/')
+',0/'+listlast(gt,'/'), gt )
| split ngt
| replace ngt replace(ngt,'/',',')
| calc thePL if(left(ngt,1)!=left(gt,1) or right(ngt,1)!=right(gt,1),listnth(pl,1,',') /* double heterozygous */
        +','+listnth(pl,int(int(right(gt,1))*(int(right(gt,1))+1)/2+int(left(gt,1))+1),',')
        +','+listnth(pl,int(int(right(ngt,1))*(int(right(ngt,1))+1)/2+int(right(ngt,1))+1),','),
        if(left(gt,1)='0',  /* heterozygous */
        listnth(pl,1,',')
        +','+listnth(pl,int(int(right(gt,1))*(int(right(gt,1))+1)/2+1),',')
        +','+listnth(pl,int(int(right(gt,1))*(int(right(gt,1))+1)/2+int(right(gt,1))+1),',')
        ,
        listnth(pl,1,',') /* homozygous */
        +','+listnth(pl,int(int(right(gt,1))*(int(right(gt,1))+1)/2+1),',')
        +','+listnth(pl,int(int(right(gt,1))*(int(right(gt,1))+1)/2+int(right(gt,1))+1),','))
        )
| calc altNum listmap(ALT,'i')
| calc theALT = listzipfilter(ALT,altnum,'listhasany(ngt,x)')
| where thealt != '<NON_REF>' and nGT != '0,0'
| calc theAD = listzipfilter(AD,ref+','+altNum,'i=1 or listhasany(ngt,x)')
| calc theMLEAC = listzipfilter(MLEAC,altNum,'listhasany(ngt,x)')
| calc theMLEAF = listzipfilter(MLEAF,altNum,'listhasany(ngt,x)')
| calc Depth = max(int(listnumsum(AD)),1)
| calc AC = listsize(AD)-if(contains(alt,'<NON_REF>'),2,1)
| calc CallRatio = form(float(listfilter(AD,'listtail(ngt)=str(i-1)'))/Depth,4,3)
| calc CallCopies if(left(ngt,1)=right(ngt,1),2,1)
/* ClippingRankSum not used */
| select 1,2,REF,theAlt,CallCopies,nGT,AC,Depth,CallRatio,GL,qual,filter,theAD,GQ,thePL,BaseQRankSum,DP
,ExcessHet,MQRankSum,RAW_MQ,ReadPosRankSum,SB,theMLEAC,theMLEAF
| rename the(.*) #{1}
| rename ngt GT
| replace GT replace(GT,',','/')
| replace PL if(AC > 1 and GL < int(listnummin(listfilter(PL,'x!="0"')))
,GL+','+listfilter(PL,'i>1'),pl) /* because we move from multi-allelic to biallelic */
| colsplit SB 4 FS | hide SB
| where not(contains(chrom,'random'))
| calc VCFpos pos
| where upper(ref) = upper(refbases(chrom,pos,pos+len(ref)-1))
| varnorm -left ref alt
| rownum | atmin 1 rownum -gc #3,#4 | hide rownum
);
```

```
create #write1# = gor [#biallelevars#]
| write #biallelefile#;

create #segcov# = pgor <(gor #filename# | #filter_chrom#
| calc depth int(vcfformattag(format,#10,'DP'))
| where depth != 'NOT_FOUND'
| calc bpEnd,bpStart int(if(alt='<NON_REF>',if(TAG(info,'END',';')!='NOT_FOUND',TAG(info,'END',';'),pos)
                                         ,str(pos+len(ref)-1) ) ) , pos-1
| select 1,bpstart,bpend,depth
| calc rd int(if(depth > 50,floor(depth/25.0)*25
                                          ,if(depth > 30,floor(depth/10.0)*10.0
                                          ,if(depth > 10, floor(depth/5.0)*5
                                          ,if(depth>5,floor(depth/2.0)*2.0
                                          ,depth)))))
| where rd > 0
| segspan -gc rd -maxseg 10000
| rename rd Depth
| hide segCount
);

create #write2# = gor [#segcov#]
| write -c #segcovfile#;

create #lowcov# = pgor [#segcov#]
| select 1-3,depth
| where depth > 0
| replace depth if(depth>10,15,depth)
| segspan -gc depth -maxseg 10000
| hide segCount;

create #write3# = gor [#lowcov#]
| write -c #lowcovfile#;

create #write4# = gor [#lowcov#]
| tee >(where depth >= 4 | segspan -maxseg 10000 | hide segCount | write -c #goodcov4# )
| tee >(where depth >= 6 | segspan -maxseg 10000 | hide segCount | write -c #goodcov6# )
| tee >(where depth >= 8 | segspan -maxseg 10000 | hide segCount | write -c #goodcov8# )
| where depth >= 10
| segspan -maxseg 10000
| hide segCount
| write -c #goodcov10#;


gorrows -p chr1:1-248956422
| top 1
```

**Example 2:** Bucketizing the GOR dictionary tables

```
def #path# = freezes/test/i2_37k_giab_ff_v9_full_genome/main/source/var;
def #input_bucket_size# = 500;

create #buckets# = nor -asdict #path#/biallele.gord
| rownum | calc bucket 'bucket_'+str(1+div(rownum-1,#input_bucket_size#))
| select PN,bucket;

create #bucketize1# = parallel -parts <(nor [#buckets#] | select bucket | distinct)
<(gor #path#/biallele.gord -ff <(nor [#buckets#] | where bucket = '#{col:bucket}')
   | write #path#/biallele_buckets/#{col:bucket}.gorz );

create #bucketize2# = parallel -parts <(nor [#buckets#] | select bucket | distinct)
<(gor #path#/lowcov.gord -ff <(nor [#buckets#] | where bucket = '#{col:bucket}')
   | write #path#/lowcov_buckets/#{col:bucket}.gorz );

/* Similar for the other table, e.g. goodcov and segcov */

/* For writing dictionary only with bucketized partitions
nor -asdict #path#/biallele.gord | map -c pn [#buckets#]
| group -gc bucket -lis -sc PN -len 100000
| calc file 'biallele_buckets/'+bucket+'.gorz'
| calc chr_start 'chr1' | calc bp_start 0 | calc chr_end 'chrZ' | calc bp_end 1000000000
| select file,bucket,chr_start,bp_start,chr_end,bp_end,lis_PN
| write #path#/biallele_bucket_partitions_only.gord
*/

nor -asdict #path#/biallele.gord | map -c pn [#buckets#]
| replace file file+'|biallele_buckets/'+bucket+'.gorz' | select file,PN
| write #path#/biallele_dual_partitions.gord
```

**Example 3:** Joint calling using the PRGTGEN command

```
def #biallele_dict# = #path#/biallele_dual_partitions.gord;
def #lowcov_dict# = #path#/lowcov_dual_partitions.gord;

create #part_allvars# = partgor -dict #biallele_dict# <(pgor #biallele_dict# -f #{tags} | group 1 -gc Ref,Alt -count);
create #allvars# = pgor -split 100 [#part_allvars#] | group 1 -gc Ref,Alt -sum -ic allcount | rename sum_allcount rowCount;

create #par_segs# = pgor [#allvars#] | group 10000 -count | seghist 1000000;

def #joint_call_bucket_size# = 5000;

create #jc_buckets# = nor -asdict #biallele_dict#
| rownum | calc bucket 'bucket_'+str(1+div(rownum-1,#joint_call_bucket_size#))
 | select PN,bucket;

create #joint_calls# = parallel -parts <(nor [#jc_buckets#] | select bucket | distinct)
<(pgor -split <(gor [#par_segs#]) #biallele_dict# -ff <(nor [#jc_buckets#] | where bucket = '#{col:bucket}' | select pn)
| select chrom,pos,ref,alt,PN,PL  /* PL triplet used instead of depth and callratio */
| merge <(gor [#allvars#] | select 1-4) /* Forces joint-calling at every variant observed globally */
| prgtgen -gc ref,alt -combgt -psep ',' -fpab 0.001 -fpbb 0.001 -pl pl -th 0.95 -e 0.001
  <(nor [#jc_buckets#] | where bucket = '#{col:bucket}')
  <(gor #lowcov_dict# -ff <(nor [#jc_buckets#] | where bucket = '#{col:bucket}' | select pn) )
);

create #write_genotypes# = parallel -parts <(nor [#jc_buckets#] | select bucket | distinct)
<(gor [#joint_calls#] -nf -ff <(nor [#jc_buckets#] | where bucket = '#{col:bucket}' | select pn)
  | write #path#/genotype_buckets/#{col:bucket}.gorz
);

create #write_dictionary# = nor [#jc_buckets#] | group -gc bucket -lis -sc PN
| calc file 'genotype_buckets/'+bucket+'.gorz'
| calc chr_start 'chr1' | calc bp_start 0 | calc chr_end 'chrZ' | calc bp_end 1000000000
| select file,bucket,chr_start,bp_start,chr_end,bp_end,lis_PN
| write #path#/variants.gord;

create #write_allvars# = pgor [#allvars#] | write #path#/metadata/allvars.gord;

nor [#jc_buckets#] | write #path#/buckets.tsv
```

#### **Example 4:** Inferring gender and calculating AF and HWE from genotypes

```
def #path# = The genotype freeze location;
def #buckets# = #path#/buckets.tsv;
def #hgt# = #path#/variants.gord; /* The horizontal genotypes */
def #allvars# = #path#/metadata/allvars.gord;

create #pns# = nor #buckets# | select pn;

create #x_segs# = gor -p chrX:2781480-155701382 #allvars#
| group 100000 -count | seghist 100000
| calc sp if(#2<2781480,2781480,#2) | calc ep if(#3>155701382,155701382,#3)
| select chrom,sp,ep,count | where ep-sp > 100;

create #Xpnvalue# = partgor -dict #hgt# -ff [#pns#] <(pgor -split <(gor [#x_segs#]) #hgt# -nf -f #{tags}
| csvsel -vs 1 -gc #3,#4 -tag pn -hide '0,3' #buckets# <(nor #buckets# | where pn in ( #{tags:q} ) | select pn)
| group genome -gc pn,value -count);

create #gen_gender# = nor [#Xpnvalue#] | select pn,value,allcount | group -gc pn,value -sum -ic allcount
| pivot value -v 1,2 -gc pn -e 0
| rename 1_sum_allcount hetCount
| rename 2_sum_allcount homCount
| calc pheno if(hetcount/(hetcount+homcount) < 0.5,'male','female');

create #varcount# = pgor #allvars# | group 100000 -count;
create #segs# = gor [#varcount#] | seghist 1000000;

def #diploid# = 2;

create #partaf# = partgor -dict #hgt# -ff [#pns#]
<(pgor -split <(gor [#segs#]) #allvars# | varjoin -norm -ir <(gor #hgt# -nf -f #{tags})
| csvcc -gc ref,alt -vs 1 -u 3 #buckets# <(nor [#gen_gender#] | where pn in ( #{tags:q} ) | select pn,pheno)
| pivot -gc ref,alt,gt cc -e 0 -v male,female
| pivot -gc ref,alt gt -v 0,1,2,3 -e 0
| prefix alt[+1]- x);

create #af# = pgor [#partaf#]
| group 1 -gc ref,alt -sum -ic x_*
| rename sum_(.*) #{1}
| calc x_AN_auto #diploid# * (x_0_female_gtcount + x_1_female_gtcount + x_2_female_gtcount
        + x_0_male_gtcount + x_1_male_gtcount + x_2_male_gtcount)
| calc x_AN_chrX #diploid# * (x_0_female_gtcount + x_1_female_gtcount + x_2_female_gtcount)
        + x_0_male_gtcount + x_1_male_gtcount + x_2_male_gtcount
| calc x_AN_chrYM x_0_female_gtcount + x_1_female_gtcount + x_2_female_gtcount
```

```
        + x_0_male_gtcount + x_1_male_gtcount + x_2_male_gtcount
| calc x_AC_auto x_1_female_gtcount + x_1_male_gtcount + #diploid# * (x_2_female_gtcount + x_2_male_gtcount)
| calc x_AC_chrX x_1_female_gtcount + x_1_male_gtcount + #diploid# * (x_2_female_gtcount) + x_2_male_gtcount
| calc x_AC_chrYM x_1_female_gtcount + x_1_male_gtcount + x_2_female_gtcount + x_2_male_gtcount
| calc AN if(chrom='chrX', x_AN_chrX, if(chrom='chrY' or chrom='chrM', x_AN_chrYM, x_AN_auto))
| calc AC if(chrom='chrX', x_AC_chrX, if(chrom='chrY' or chrom='chrM', x_AC_chrYM, x_AC_auto))
| calc x_0_gtcount x_0_female_gtcount + x_0_male_gtcount
| calc x_1_gtcount x_1_female_gtcount + x_1_male_gtcount
| calc x_2_gtcount x_2_female_gtcount + x_2_male_gtcount
| calc x_3_gtcount x_3_female_gtcount + x_3_male_gtcount
| calc AF form(AC/float(AN),6,7)
| calc conservative_AF form(max(0.0, float(AC - 2*sqrt(AC))/(AN)),6,7)
| calc pnCount x_0_gtcount+x_1_gtcount+x_2_gtcount+x_3_gtcount
| calc marker_yield form((float(x_0_gtcount)+float(x_1_gtcount)+float(x_2_gtcount))/float(pncount),6,7)

/* HWE assumes diploid, so we ignore sex */
| calc HWE_homFreq form(x_2_gtcount/float(PNcount),6,7)
| calc hetFreq form(x_1_gtcount/float(pncount),6,7)
| calc HWE_alleleFreq form((2*x_2_gtcount+x_1_gtcount)/float(2*pncount),6,7)
| calc pA = (2*x_2_gtcount+x_1_gtcount)/float(2*PNcount)
| calc pB = 1.0 - pA
| calc mAA = pA * pA * PNcount
| calc mAB = 2 * pA * pB * PNcount
| calc mBB = pB * pB * PNcount
| calc HWE_chisq = float(form(sqr(max(abs(x_2_gtcount-mAA)-0.5,0.0))/mAA
                 + sqr(max(abs(x_1_gtcount-mAB)-0.5,0.0))/mAB + sqr(max(abs(PNcount-x_1_gtcount-x_2_gtcount-mBB)-0.5,0.0))/mBB ,5,5))
| calc HWE_pVal = chisquarecompl(1,HWE_chisq)
| calc HWE_logPval = float(form(-log(HWE_pVal),3,2))
| calc varCount x_1_gtcount + x_2_gtcount
| calc hetCount x_1_gtcount
| calc homCount x_2_gtcount
| hide pA-mBB
| hide x_*;

pgor [#af#] | write #path#/metadata/AF.gord
```

**Example 5:** Calculating genome region based metrics from the AF table

```
def #af# = #path#/metadata/AF.gord;

create #allsingle# = pgor #af#
| calc single if(ac=1,1,0)
| group 100 -steps 5 -count -sum -ic single
| where sum_single = allcount and allcount > 2
| segspan -maxseg 10000;

create #highpoly# = pgor #af#
| group 1 -gc ref -count
| where allcount > 1
| group 100 -steps 5 -count
| where allcount > 4
| segspan -maxseg 10000;

create #lowyield# = pgor #af#
| group 100 -steps 5 -avg -fc marker_yield
| where avg_marker_yield < 0.99
| segspan -maxseg 10000;

create #repeatSingletons# = pgor #af#
| where len(#3)!=len(#4)
| select 1-4
| calc oldpos pos-1
| varnorm -right #3 #4
| select 1,oldpos,pos
| rename #2 varstart
| rename #3 varStop
| sort 10000
| where #3-#2>0
| calc size #3-#2
| join -segsnp -ic <(gor #af# | where ac < 2)
| calc overlapRatio float(form(float(overlapcount/size),3,2))
| where overlapratio > 2
| segspan -maxseg 10000;

create #repeatRegion# = pgor #af#
| where len(#3)!=len(#4)
| select 1-4
| calc oldpos pos-1
| varnorm -right #3 #4
| select 1,oldpos,pos
| rename #2 varstart
```

```
| rename #3 varStop
| where #3-#2 > 10
| sort 10000
| segspan -maxseg 10000;

create #region_qc# = pgor #af#
| select 1-4
| join -snpseg -maxseg 10000 -ic [#allsingle#]
| join -snpseg -maxseg 10000 -ic [#highpoly#]
| join -snpseg -maxseg 10000 -ic [#lowyield#]
| join -snpseg -maxseg 10000 -ic [#repeatSingletons#]
| join -snpseg -maxseg 10000 -ic [#repeatRegion#]
| replace overlapcount* if(int(#rc)>0,1,0)
| rename overlapcount allsingle
| rename overlapcountx highpoly
| rename overlapcountxx lowyield
| rename overlapcountxxx repeatSingletons
| rename overlapcountxxxx repeatRegion
;
```

**Example 6:** Finding total sequence read depth and the frequency of poor coverage

```
def #segcov# = #path#/segcov.gord;
def #goodcov4# = #path#/goodcov_4.gord;
def #goodcov6# = #path#/goodcov_6.gord;
def #goodcov8# = #path#/goodcov_8.gord;
def #goodcov10# = #path#/goodcov_10.gord;

def #numSamplesPlusOne# = 37023; /* This number should be one larger than the samples in joint-calling */

create #zeros# = gorrows -p chr1:1-2
| calc newchrom 'chr1,chr10,chr11,chr12,chr13,chr14,chr15,chr16,chr17,chr18,chr19,'
               +'chr2,chr20,chr21,chr22,chr3,chr4,chr5,chr6,chr7,chr8,chr9,chrM,chrX,chrY'
| split newchrom | select newchrom,pos
| group chrom -count
| segspan -maxseg 10000
| segproj -maxseg 100000;

create #p4# = partgor -dict #goodcov4# -partsize 5000 <(pgor #goodcov4# -f #{tags}  | segproj -maxseg 10000 );
create #lowCovQC4# = pgor <(gor [#p4#] | merge [#zeros#] | segproj -maxseg 10000 -sumcol segcount
| replace segCount #numSamplesPlusOne# - segCount | rename segCount lowCov4);

create #p6# = partgor -dict #goodcov6# -partsize 5000 <(pgor #goodcov6# -f #{tags}  | segproj -maxseg 10000 );
create #lowCovQC6# = pgor <(gor [#p6#] | merge [#zeros#] | segproj -maxseg 10000 -sumcol segcount
| replace segCount #numSamplesPlusOne# - segCount | rename segCount lowCov6);

create #p8# = partgor -dict #goodcov8# -partsize 5000 <(pgor #goodcov8# -f #{tags}  | segproj -maxseg 10000 );
create #lowCovQC8# = pgor <(gor [#p8#] | merge [#zeros#] | segproj -maxseg 10000 -sumcol segcount
| replace segCount #numSamplesPlusOne# - segCount | rename segCount lowCov8);

create #p10# = partgor -dict #goodcov10# -partsize 5000 <(pgor #goodcov10# -f #{tags}  | segproj -maxseg 10000 );
create #lowCovQC10# = pgor <(gor [#p10#] | merge [#zeros#] | segproj -maxseg 10000 -sumcol segcount
| replace segCount #numSamplesPlusOne# - segCount | rename segCount lowCov10);

create #depth_parts# = partgor -dict #segcov# -partscale 4  <(pgor -split 50000000:10000 #segcov# -f #{tags}
| segproj -maxseg 10000 -sumcol depth );

create #depth_segs# = pgor -split 10000000:10000 [#depth_parts#] | segproj -maxseg 10000 -sumcol segcount
| rename segcount Depth | calc nbpStart,nbpStop max(#{BPSTART}+9999,#2),min(#3,#{BPSTOP}-10000)
| select #1,nbpStart,nbpStop,depth
| rename #2 bpStart | rename #3 bpStop
| where #3 > #2;
```

**Example 7:** Aggregating variant attributes

```
def #biallele_dict# = #path#/biallele_dual_partitions.gord;
def #allvars# = #path#/metadata/allvars.gord;

create #segcount# = pgor -split 200 #allvars#  | group 100000 -sum -ic rowCount;

create #segs# = gor [#segcount#]  | seghist 300000000; /* makes each job about 3hrs  */

create #varqc# = pgor -split <(gor [#segs#]) #biallele_dict#
| calc GQLO2 if(GQ<2,1,0)
| calc GQLO5 if(GQ<5,1,0)
| calc GQLO10 if(GQ<10,1,0)
| calc GQLO20 if(GQ<20,1,0)
| calc GQLO30 if(GQ<30,1,0)
| calc GQGRE2 if(GQ>=2,1,0)
```

```
| calc GQGRE5 if(GQ>=5,1,0)
| calc GQGRE10 if(GQ>=10,1,0)
| calc GQGRE20 if(GQ>=20,1,0)
| calc GQGRE25 if(GQ>=25,1,0)
| calc GQGRE30 if(GQ>=30,1,0)
| calc GQGRE35 if(GQ>=35,1,0)
| calc GQGRE40 if(GQ>=40,1,0)
| calc GQGRE50 if(GQ>=50,1,0)
| calc AC2 if(AC>1,1,0)
| calc AC3 if(AC>2,1,0)
| calc AC4 if(AC>3,1,0)
| replace baseqranksum,mqranksum,readposranksum if(callratio > 0.0001 and callratio < 0.999,#rc,'NaN')
| group 1 -gc ref,alt -sum -med -ic gl,depth,DP,QUAL,GQ*,AC*,fs_* -fc baseqranksum,mqranksum,readposranksum,raw_mq -count
| rename sum_fs_1 ref_fw | rename sum_fs_2 ref_rv | rename sum_fs_3 alt_fw | rename sum_fs_4 alt_rv
| calc SOR form(ln( (ref_fw+1)*(alt_rv+1)/((ref_rv+1)*(alt_fw+1)) + ((ref_rv+1)*(alt_fw+1))/((ref_fw+1)*(alt_rv+1)) )
+ ln(min(ref_rv+1,ref_fw+1)/max(ref_rv+1,ref_fw+1)) - ln(min(alt_fw+1,alt_rv+1)/max(alt_rv+1,alt_fw+1)),4,3)
| calc FS form(-10*log(pval(int(ref_fw),int(ref_rv),int(alt_fw),int(alt_rv))),4,3)
| calc FSzero if(ref_fw = 0 or alt_fw =0 or ref_rv =0 or alt_rv = 0,1,0)
| calc MQ form(sqrt(sum_raw_mq/float(sum_depth)),4,2)
| calc QD form(sum_qual/sum_depth,4,2)
| rename allcount SamplesWithVar
| rename med_BaseQRankSum BaseQRankSum
| rename med_MQRankSum MQRankSum
| rename med_ReadPosRankSum ReadPosRankSum
| rename sum_GQLO(.*) GQLO#{1}
| rename sum_GQGRE(.*) GQGRE#{1}
| rename sum_AC(.+) AC#{1}
| select 1,2,Ref,Alt,SamplesWithVar,QD,FS,FSzero,MQ,SOR,BaseQRankSum
            ,MQRankSum,ReadPosRankSum,med_Depth,AC2,AC3,AC4
            ,GQGRE10,GQGRE2,GQGRE20,GQGRE25,GQGRE30,GQGRE35,GQGRE40,GQGRE5,GQGRE50
            ,GQLO10,GQLO2,GQLO20,GQLO30,GQLO5
;

def #af# = #path#/metadata/AF.gord;

create #normalized_varqc# = pgor -split 100 #af#
| calc inbreeding if(af=0.0 or af=1.0,0.0,1 - (hetcount)/(AN*af*(1-af)))
| prefix 5- freezeStat
| varjoin -r -rprefix region [#region_qc#]
| varjoin -norm -r -rprefix callStat [#varqc#]
| join -snpseg -r -l -e 0 -maxseg 10000 [#lowCovQC4#]
| join -snpseg -r -l -e 0 -maxseg 10000 [#lowCovQC6#]
| join -snpseg -r -l -e 0 -maxseg 10000 [#lowCovQC8#]
| join -snpseg -r -l -e 0 -maxseg 10000 [#lowCovQC10#]
| rename callStat_sum_GQ(.*) callStat_GQ#{1}
| rename callStat_sum_AC(.*) callStat_AC#{1}
| join -snpseg -maxseg 10000 -l -e 0 -r [#depth_segs#]
| rename Depth freezeStat_sum_DP
| calc freezeStat_avg_DP form(freezeStat_sum_DP/freezeStat_pnCount,4,2)
| select 1,2,ref,alt,freezeStat_AN,freezeStat_AC,freezeStat_AF,freezeStat_conservative_AF
            ,freezeStat_pnCount,freezeStat_marker_yield
            ,freezeStat_avg_DP,freezeStat_sum_DP,freezeStat_HWE_chisq,freezeStat_HWE_pVal,freezeStat_HWE_logPval
            ,freezeStat_varCount,freezeStat_hetCount,freezeStat_homCount,freezeStat_inbreeding
            ,region_allsingle,region_highpoly,region_lowyield,region_repeatSingletons,region_repeatRegion
            ,callStat_SamplesWithVar,callStat_QD,callStat_FS,callStat_FSzero,callStat_MQ,callStat_SOR,callStat_BaseQRankSum
            ,callStat_MQRankSum,callStat_ReadPosRankSum,callStat_med_Depth
            ,callStat_GQLO2,callStat_GQLO5,callStat_GQLO10,callStat_GQLO20,callStat_GQLO30
            ,callStat_GQGRE2,callStat_GQGRE5,callStat_GQGRE10,callStat_GQGRE20,callStat_GQGRE25,callStat_GQGRE30
            ,callStat_GQGRE35,callStat_GQGRE40,callStat_GQGRE50,callStat_AC2,callStat_AC3,callStat_AC4
            ,lowCov4,lowCov6,lowCov8,lowCov10
| prefix lowCov4-lowCov10 cov
| replace callstat_GQ* form(float(#rc)/callstat_SamplesWithVar,5,5)
| replace callstat_AC* form(float(#rc)/callstat_SamplesWithVar,5,5)
| replace cov_* form(float(#rc)/freezestat_pncount,5,5)
| replace freezeStat_marker_yield,freezeStat_inbreeding,freezeStat_HWE_logPval,freezeStat_HWE_chisq form(float(#rc),4,6)
| join -snpsnp -l -e 0 -rprefix Ill
        <(gor user_data/hakon/freezes/test/i2_37k_ff_v6/refbaseFullQC.gord
          | select 1,2,avg_fbQ20,avDepth,avg_mapQ
         | rename avDepth avg_Depth
         | calc seqBad 3- (if(avg_mapQ>=50,1,0)
                                 + if(chrom!='chrX' and avg_Depth>=0.75 and avg_Depth<=1.25
                                        or chrom='chrX' and avg_Depth>=0.58 and avg_Depth<=0.97 ,1,0) + if(avg_fbQ20>=0.90,1,0)) )
| join -snpseg -ic user_data/hakon/freezes/test/i2_37k_ff_v6/GRCh38_alldifficultregions.gorz
| rename overlapcount GIAB_allDiffRegions
;

pgor -split 100 [#normalized_varqc#] | write #path#/metadata/variantQC.gord
```

**Example 8:** Calculating regional alignment quality metrics

```
create #pns# = nor -h user_data/hakon/freezeQC.tsv
| where avgdepth > 30 and avgdepth<30.15
```

```
| select pn | inset -c pn <(nor -asdict source/bam/bam.gord | select #2)
| top 32;

create xxx = partgor -dict source/bam/bam.gord -ff [#pns#] -partsize 1 <(pgor <(gor source/bam/bam.gord -f #{tags}
| calc Map50 if(mapq>=50,'H','L')
| pileup -gc Map50 -bq 0 -depth
| hide refbase
| pivot map50 -v H,L -e 0
| join -snpsnp -l -e 0 <(gor source/bam/bam.gord -f #{tags}
| calc Map50 if(mapq>=50,'H','L')
| pileup -gc Map50 -bq 20 -depth
| rename depth Depth20
| hide refbase
| pivot map50 -v H,L -e 0)
)
| calc fMap50 form(h_depth/(h_depth+l_depth),3,3)
| calc fbQ20 form((h_depth20+l_depth20)/(h_depth+l_depth),3,3)
| calc Depth form((H_Depth+L_Depth)/38.34,3,3)
| select chrom,pos,depth,fmap50,fbq20
);

create xxx2 = partgor -dict source/bam/bam.gord -ff [#pns#] -partsize 1 <(pgor source/bam/bam.gord -f #{tags}
| select 1-3,mapq | join -segsnp <(gor config/chromSeqhg38)
| select #1,posx,mapq
| sort 5000
| group 1 -avg -ic mapq
| replace avg_mapq form(float(avg_mapq),3,2)
)
;

create yyy = pgor -split 100 [xxx] | group 1 -avg -std -ic depth -fc fmap50,fbq20 -count
| replace std_* float(#rc)/sqrt(allcount)
| replace 4- form(float(#rc),3,3)
| calc avDepth if(allcount<32,form(float(avg_depth)*allcount/32,3,3),avg_depth)
| write user_data/hakon/freezes/test/i2_37k_ff_v6/refbaseQC.gord;

create yyy2 = pgor -split 100 [xxx2] | group 1 -avg -std -ic depth -fc avg_mapq -count
| replace std_* float(#rc)/sqrt(allcount)
| replace 4- form(float(#rc),3,3)
| write user_data/hakon/freezes/test/i2_37k_ff_v6/refbaseMapQ.gord;

create zzz = pgor -split 100 user_data/hakon/freezes/test/i2_37k_ff_v6/refbaseQC.gord
| join -snpsnp -l -e 0 -r <(gor user_data/hakon/freezes/test/i2_37k_ff_v6/refbaseMapQ.gord )
| write user_data/hakon/freezes/test/i2_37k_ff_v6/refbaseFullQC.gord;

gor #genes# | top 1
```

#### **Example 9:** Converting GIAB VCF and BED files to GOR

```
def #GIABpath# = freezes/test/i2_37k_giab_ff_v8/main/source/var/vcf_giab;

create #vcfHG001# = gor #GIABpath#/HG001_GRCh38_1_22_v4.2.1_benchmark.vcf.gz
| rename #10 Data
| hide id
| calc GT vcfformattag(format,data,'GT')
| columnsort chrom,pos,ref,alt,gt
| sort genome
| calc sample 'HG001';

create #vcfNA12878# = gor [#vcfHG001#] | replace sample 'NA12878';

create #vcfHG002# = gor #GIABpath#/HG002_GRCh38_1_22_v4.2.1_benchmark.vcf.gz
| rename #10 Data
| hide id
| calc GT vcfformattag(format,data,'GT')
| columnsort chrom,pos,ref,alt,gt
| sort genome
| calc sample 'HG002';

create #vcfHG003# = gor #GIABpath#/HG003_GRCh38_1_22_v4.2.1_benchmark.vcf.gz
| rename #10 Data
| hide id
| calc GT vcfformattag(format,data,'GT')
| columnsort chrom,pos,ref,alt,gt
| sort genome
| calc sample 'HG003';

create #vcfHG004# = gor #GIABpath#/HG004_GRCh38_1_22_v4.2.1_benchmark.vcf.gz
| rename #10 Data
| hide id
| calc GT vcfformattag(format,data,'GT')
| columnsort chrom,pos,ref,alt,gt
```

```
| sort genome
| calc sample 'HG004';

create #vcfHG005# = gor #GIABpath#/HG005_GRCh38_1_22_v4.2.1_benchmark.vcf.gz
| rename #10 Data
| hide id
| calc GT vcfformattag(format,data,'GT')
| columnsort chrom,pos,ref,alt,gt
| sort genome
| calc sample 'HG005';

create #vcfHG006# = gor #GIABpath#/HG006_GRCh38_1_22_v4.2.1_benchmark.vcf.gz
| rename #10 Data
| hide id
| calc GT vcfformattag(format,data,'GT')
| columnsort chrom,pos,ref,alt,gt
| sort genome
| calc sample 'HG006';

create #vcfHG007# = gor #GIABpath#/HG007_GRCh38_1_22_v4.2.1_benchmark.vcf.gz
| rename #10 Data
| hide id
| calc GT vcfformattag(format,data,'GT')
| columnsort chrom,pos,ref,alt,gt
| sort genome
| calc sample 'HG007';

create #bedHG001# = gor <(nor #GIABpath#/HG001_GRCh38_1_22_v4.2.1_benchmark.bed
| rename #1 Chrom
| rename #2 bpStart
| rename #3 bpStop)
| sort genome;

create #bedHG002# = gor <(nor #GIABpath#/HG002_GRCh38_1_22_v4.2.1_benchmark_noinconsistent.bed
| rename #1 Chrom
| rename #2 bpStart
| rename #3 bpStop)
| sort genome;

create #bedHG003# = gor <(nor #GIABpath#/HG003_GRCh38_1_22_v4.2.1_benchmark_noinconsistent.bed
| rename #1 Chrom
| rename #2 bpStart
| rename #3 bpStop)
| sort genome;

create #bedHG004# = gor <(nor #GIABpath#/HG004_GRCh38_1_22_v4.2.1_benchmark_noinconsistent.bed
| rename #1 Chrom
| rename #2 bpStart
| rename #3 bpStop)
| sort genome;

create #bedHG005# = gor <(nor #GIABpath#/HG005_GRCh38_1_22_v4.2.1_benchmark.bed
| rename #1 Chrom
| rename #2 bpStart
| rename #3 bpStop)
| sort genome;

create #bedHG006# = gor <(nor #GIABpath#/HG006_GRCh38_1_22_v4.2.1_benchmark.bed
| rename #1 Chrom
| rename #2 bpStart
| rename #3 bpStop)
| sort genome;

create #bedHG007# = gor <(nor #GIABpath#/HG007_GRCh38_1_22_v4.2.1_benchmark.bed
| rename #1 Chrom
| rename #2 bpStart
| rename #3 bpStop)
| sort genome;

create #goodsegs# = gor [#bedHG001#] | calc sample 'HG001'
| merge <(gor [#bedHG001#] | calc sample 'NA12878')
| merge <(gor [#bedHG005#] | calc sample 'HG002')
| merge <(gor [#bedHG005#] | calc sample 'HG003')
| merge <(gor [#bedHG005#] | calc sample 'HG004')
| merge <(gor [#bedHG005#] | calc sample 'HG005')
| merge <(gor [#bedHG006#] | calc sample 'HG006')
| merge <(gor [#bedHG007#] | calc sample 'HG007');

create #vcfs# = gor [#vcfHG001#] [#vcfNA12878#] [#vcfHG002#]
                                  [#vcfHG003#] [#vcfHG004#] [#vcfHG005#] [#vcfHG006#] [#vcfHG007#]
| replace gt replace(gt,'|','/')
| where gt != '0/0'
| calc ngt if(listfirst(gt,'/') != '0' and listfirst(gt,'/') != listlast(gt,'/')
                   ,'0/'+listfirst(gt,'/')+',0/'+listlast(gt,'/'), gt )
```

```
| split ngt
| replace ngt replace(ngt,'/',',')
| calc altNum listmap(ALT,'i')
| calc theALT = listzipfilter(ALT,altnum,'listhasany(ngt,x)')
| replace theALT if(#rc='*','',#rc)
| where thealt != '<NON_REF>' and nGT != '0,0'
| calc CallCopies if(left(ngt,1)=right(ngt,1),2,1)
| hide alt
| rename thealt Alt
| select 1,2,ref,alt,callcopies,sample;
```

##### **Example 10:** Calculating VCF variant count and BED coverage fraction

```
nor <(gor [#goodsegs#] | calc size bpstop-bpstart | group chrom -sum -ic size -gc sample | calc chr_size #3-#2)
| group -gc sample -sum -ic sum_size,chr_size | calc fraction sum_sum_size/sum_chr_size | select sample,fraction
| map -c sample <(nor [#vcfs#] | group -gc sample -count | rename allcount varCount)
```

##### **Example 11:** Generating balanced training data for XGBoost

```
/* Use definitions from Ex.9 */

def #pns# = 'HG001','NA12878','HG002','HG003','HG004','HG006','HG007';

def #biallele_path# = freezes/test/i2_37k_giab_ff_v9_full_genome/main;
def #GATKfreeze# = freezes/test/i2_37k_giab_new_sention/main;
def #GORfreeze# = freezes/test/i2_37k_giab_ff_v9_full_genome/main;

create #vars_per_pn# = pgor  <(gor [#vcfs#] | select 1,2,ref,alt,sample | rename sample PN
| merge <(gor #biallele_path#/source/var/biallele.gord -f #pns# | select 1-4,PN | rename #3 Ref | rename #4 Alt))
| group 1 -gc ref,alt,pn;

create #GATKparsegs# = pgor  #GATKfreeze#/metadata/AF.gorz | group 100000 -count | seghist 1000000;

create #GATKfreezevars# = pgor -split <(gor [#GATKparsegs#]) #GATKfreeze#/variants.gord -f #pns# -nf
| csvsel -gc ref,alt -vs 1 -u 3 #GATKfreeze#/buckets.tsv
             <(nor #GATKfreeze#/buckets.tsv | where pn in ( #pns# ) | select pn) -tag PN -hide 0;

create #GATKvars_per_pn# = pgor <(gor [#vcfs#] | select 1,2,ref,alt,sample | rename sample PN
| merge <(gor [#GATKfreezevars#] | select 1-4,PN | rename #3 Ref | rename #4 Alt)) | group 1 -gc ref,alt,pn;

create #GORparsegs# = pgor  #GORfreeze#/metadata/AF.gorz | group 100000 -count | seghist 1000000;

create #GORfreezevars# = pgor -split <(gor [#GORparsegs#]) #GORfreeze#/variants.gord -f #pns# -nf
| csvsel -gc ref,alt -vs 1 -u 3 #GORfreeze#/buckets.tsv
             <(nor #GORfreeze#/buckets.tsv | where pn in ( #pns# ) | select pn) -tag PN -hide 0;

create #GORvars_per_pn# = pgor  <(gor [#vcfs#] | select 1,2,ref,alt,sample | rename sample PN
| merge <(gor [#GORfreezevars#] | select 1-4,PN | rename #3 Ref | rename #4 Alt)) | group 1 -gc ref,alt,pn;

create #posnegvars# = pgor [#GORvars_per_pn#]
| where pn in ( #pns# )
| join -snpseg -i -xl pn -xr sample [#goodsegs#]
| varjoin -r -l -xl pn -xr sample [#vcfs#]
| varjoin -r -l -xl pn -xr pn <(gor [#GORfreezevars#] | rename value Callcopies )
| select 1,2,ref,alt,pn,callcopies,callcopiesx
| calc TP if(callcopies != '' and callcopies = callcopiesx,1,0)
| calc FN if(callcopies != '' and callcopies != callcopiesx,1,0)
| calc FP if(callcopies = '' and callcopiesx != '',1,0)
| group 1 -gc ref,alt -sum -ic tp,fn,fp
| calc tp if(sum_tp>0,1,0)
| calc fn if(sum_fn>0,1,0)
| calc fp if(sum_fp>0,1,0)
| calc posneg if(tp=1 and fn=0 and fp=0,'pos',if(tp=0 and (fn=1 or fp=1),'neg','other'))
| where posneg in ('pos','neg')
| select 1-4,posneg;

create #training# = gor [#posnegvars#] | where chrom != 'chr2'
| where posneg = 'neg' or random()<0.1;

create #testing# = gor [#posnegvars#] | varjoin -n [#training#];

gor [#training#]
| calc snpInDel if(len(ref)=len(alt),'SNP','InDel')
| varjoin -r <(gor #path#/metadata/variantQC.gord ) /* see query in Ex.7 */
| write #path#/training.gorz
```

##### **Example 12:** GIAB concordance analysis

```
/* Use definitions from Ex.9 and Ex.11 */
```

```
create #comp_bialleele# = gor -p chr22 [#vars_per_pn#]
| where pn in ( #pns# )
| join -snpseg -i -xl pn -xr sample [#goodsegs#]
| varjoin -ic <(gor user_data/hakon/freezes/test/i2_37k_giab_ff_v9_full_genome/v2/pred_thin.gorz | where prediction = 1)
| calc good if(overlapcount > 0,1,0)
| varjoin -r -l -xl pn -xr sample [#vcfs#]
| varjoin -r -l -xl pn -xr pn <(gor #biallele_path#/source/var/biallele.gord -f #pns# )
| select 1,2,ref,alt,pn,callcopies,callcopiesx,good
| join -snpseg -l -e 0 -xl pn -xr pn -maxseg 10000 -r <(gor #biallele_path#/source/cov/segcov.gord -f #pns# )
| hide pnx
| calc snp if(len(ref)=len(alt),1,0)
| replace depth if(depth < 8,0,1)
| group genome -gc pn,callcopies,callcopiesx,depth,snp,good -count;

create #GORcomp_freeze# = gor -p chr22 [#GORvars_per_pn#]
| where pn in ( #pns# )
| join -snpseg -i -xl pn -xr sample [#goodsegs#]
| varjoin -ic <(gor user_data/hakon/freezes/test/i2_37k_giab_ff_v9_full_genome/v2/pred_thin.gorz | where prediction = 1)
| calc good if(overlapcount > 0,1,0)
| varjoin -r -l -xl pn -xr sample [#vcfs#]
| varjoin -r -l -xl pn -xr pn <(gor [#GORfreezevars#] | rename value Callcopies )
| select 1,2,ref,alt,pn,callcopies,callcopiesx,good
| join -snpseg -l -e 0 -xl pn -xr pn -maxseg 10000 -r <(gor #GORfreeze#/source/cov/segcov.gord -f #pns# )
| hide pnx
| calc snp if(len(ref)=len(alt),1,0)
| replace depth if(depth < 8,0,1)
| group chrom -gc pn,callcopies,callcopiesx,depth,snp,good -count;

create #GATKcomp_freeze# = gor -p chr22 [#GATKvars_per_pn#]
| where pn in ( #pns# )
| join -snpseg -i -xl pn -xr sample [#goodsegs#]
| varjoin -ic <(gor user_data/hakon/freezes/test/i2_37k_giab_ff_v9_full_genome/v2/pred_thin.gorz | where prediction = 1)
| calc good if(overlapcount > 0,1,0)
| varjoin -r -l -xl pn -xr sample [#vcfs#]
| varjoin -r -l -xl pn -xr pn <(gor [#GATKfreezevars#] | rename value Callcopies )
| select 1,2,ref,alt,pn,callcopies,callcopiesx,good
| join -snpseg -l -e 0 -xl pn -xr pn -maxseg 10000 -r <(gor #biallele_path#/source/cov/segcov.gord -f #pns# )
| hide pnx
| calc snp if(len(ref)=len(alt),1,0)
| replace depth if(depth < 8,0,1)
| group genome -gc pn,callcopies,callcopiesx,depth,snp,good -count;

create #bialleleQC# = nor [#comp_bialleele#]
| #depthfilter#
| #snpfilter#
| #goodfilter#
| calc TP if(callcopies != '' and callcopies = callcopiesx,allcount,0)
| calc FN if(callcopies != '' and callcopies != callcopiesx,allcount,0)
| calc FP if(callcopies = '' and callcopiesx != '',allcount,0)
##collapsePNs##
| group -gc pn -sum -ic TP,FN,FP
| rename sum_(.*) #{1}
| calc TPR TP/(TP+FN) /* Sensitivity */
| calc PPV TP/(TP+FP) /* Precision */
| calc F1 2*TPR*PPV/(TPR+PPV)
| replace TPR- form(float(#rc),5,4)
| calc VARS TP+FN
| prefix 2- biallele
;

create #GORfreezeQC# = nor [#GORcomp_freeze#]
| #depthfilter#
| #snpfilter#
| #goodfilter#
| where callcopiesx != '3'
| calc TP if(callcopies != '' and callcopies = callcopiesx,allcount,0)
| calc FN if(callcopies != '' and callcopies != callcopiesx,allcount,0)
| calc FP if(callcopies = '' and callcopiesx != '',allcount,0)
##collapsePNs##
| group -gc pn -sum -ic TP,FN,FP
| rename sum_(.*) #{1}
| calc TPR TP/(TP+FN) /* Sensitivity */
| calc PPV TP/(TP+FP) /* Precision */
| calc F1 2*TPR*PPV/(TPR+PPV)
| replace TPR- form(float(#rc),5,4)
| calc VARS TP+FN
| prefix 2- Freeze;

create #GATKfreezeQC# = nor [#GATKcomp_freeze#]
| #depthfilter#
| #snpfilter#
| #goodfilter#
| where callcopiesx != '3'
```

```
| calc TP if(callcopies != '' and callcopies = callcopiesx,allcount,0)
| calc FN if(callcopies != '' and callcopies != callcopiesx,allcount,0)
| calc FP if(callcopies = '' and callcopiesx != '',allcount,0)
##collapsePNs##
| group -gc pn -sum -ic TP,FN,FP
| rename sum_(.*) #{1}
| calc TPR TP/(TP+FN) /* Sensitivity */
| calc PPV TP/(TP+FP) /* Precision */
| calc F1 2*TPR*PPV/(TPR+PPV)
| replace TPR- form(float(#rc),5,4)
| calc VARS TP+FN
| prefix 2- Freeze;

def #snpfilter# = where snp = 1;
def #depthfilter# = where depth >= 0;
def #goodfilter# = where good >= 1;
def ##collapsePNs## = /* | replace pn 'all'  */ ;

create #GOR# = nor [#GORfreezeQC#]
| map -c PN <(nor [#bialleleQC#])
| rename PN Sample
| rename biallele_vars Vars
| select  Sample,VARS,Freeze_TPR,biallele_TPR,Freeze_PPV,biallele_PPV,Freeze_F1,biallele_F1;

create #GATK# = nor [#GATKfreezeQC#]
| map -c PN <(nor [#bialleleQC#])
| rename PN Sample
| rename biallele_vars Vars
| select  Sample,VARS,Freeze_TPR,biallele_TPR,Freeze_PPV,biallele_PPV,Freeze_F1,biallele_F1;

nor [#GOR#] | rename freeze_(.*) GOR_#{1}
| map -c #1 <(nor [#GATK#] | hide vars,biallele_* | rename freeze_(.*) GATK_#{1})
| select  Sample,Vars,biallele_TPR,GATK_TPR,GOR_TPR,biallele_PPV,GATK_PPV,GOR_PPV,biallele_F1,GATK_F1,GOR_F1
```

#### **Example 13:** Training XGBoost in SparkGOR

```
# Start pyspark

pyspark --repositories https://s3-us-west-2.amazonaws.com/xgboost-maven-repo/snapshot/
              --packages "ml.dmlc:xgboost4j_2.12:1.7.0-SNAPSHOT,org.gorpipe:gor-spark:4.2.9"
              --exclude-packages "com.google.inject:guice,io.netty:netty-codec-http"

# Then run this SparkGOR script

from pyspark.sql import SparkSession
import gor_pyspark
spark = SparkSession.builder.getOrCreate()
sgs = spark.createGorSession()

queryrest = """
| rename posneg label
| replace label if(label='neg',0,1)
| replace snpInDel if(snpInDel='SNP',0,1)
| replace callStat_FS if(callStat_FS='Infinity',2147483647,callStat_FS)
| replace callStat_QD if(callStat_QD='Infinity',2147483647,callStat_QD)
| replace callStat_BaseQRankSum if(callStat_BaseQRankSum='Infinity',2147483647,callStat_BaseQRankSum)
| replace callStat_MQRankSum if(callStat_MQRankSum='Infinity',2147483647,callStat_MQRankSum)
| replace callStat_ReadPosRankSum if(callStat_ReadPosRankSum='Infinity',2147483647,callStat_ReadPosRankSum)
| replace freezeStat_HWE_chisq if(freezeStat_HWE_chisq='Infinity',2147483647,freezeStat_HWE_chisq)
| replace freezeStat_HWE_logPval if(freezeStat_HWE_logPval='Infinity',2147483647,freezeStat_HWE_logPval)
| where not(freezeStat_HWE_chisq='NaN')
| where not(freezeStat_HWE_logPval='NaN')
| cols2list -gc REF,ALT,label snpInDel,freezeStat_sum_DP,freezeStat_AC,freezeStat_AF,freezeStat_marker_yield,freezeStat_HWE_chisq
,freezeStat_HWE_logPval,freezeStat_varCount,freezeStat_hetCount,freezeStat_homCount,freezeStat_inbreeding
,region_allsingle,region_highpoly,region_lowyield,region_repeatSingletons,region_repeatRegion,callStat_SamplesWithVar
,callStat_QD,callStat_FS,callStat_SOR,callStat_MQ,callStat_med_Depth,callStat_BaseQRankSum,callStat_MQRankSum
,callStat_ReadPosRankSum,callStat_GQLO2,callStat_GQLO5,callStat_GQLO10,callStat_GQLO20,callStat_GQLO30
,callStat_GQGRE2,callStat_GQGRE5,callStat_GQGRE10,callStat_GQGRE20,callStat_GQGRE25,callStat_GQGRE30
,callStat_GQGRE35,callStat_GQGRE40,callStat_GQGRE50,callStat_AC2,callStat_AC3,callStat_AC4,cov_lowCov4
,cov_lowCov6,cov_lowCov8,cov_lowCov10,Ill_avg_fbQ20,Ill_avg_Depth,Ill_avg_mapQ,Ill_seqBad features
"""
trainquery = f"pgor #path#/metadata/variantQC.gord | calc snpInDel if(len(ref)=len(alt),'SNP','InDel'} {queryrest}"


ddlschema = "chrom string,pos int,ref string,alt string,label int,features string"

traindf = sgs.pydataframe(trainquery, ddlschema)

# Register udfs
from pyspark.sql.types import *
spark.udf.registerJavaFunction("strlisttoarray","org.gorpipe.spark.udfs.CommaToDoubleArray",ArrayType(DoubleType()))
```

```
# Change gor string list to spark ml vector
from pyspark.sql.functions import *
trainaf = traindf.withColumn("features",expr("strlisttoarray(features)"))

from xgboost.spark import SparkXGBClassifier
spark_reg_estimator = SparkXGBClassifier(features_col="features", \
                                         label_col="label", \
                                         use_gpu=False, \
                                         num_workers=int(6), \
                                         evalmetric="logloss", \
                                         learning_rate=float(0.1), \
                                         num_round=500, \
                                         n_thread=2, \
                                         max_depth=8, \
                                         subsample=0.5)
model = spark_reg_estimator.fit(trainaf)

# model.save(f"{projectdir}/{resultdir}/mymodel")

testquery = f"pgor #path#/training.gorz | calc snpInDel if(len(ref)=len(alt),'SNP','InDel') {queryrest}"
testdf = sgs.pydataframe(testquery, ddlschema)
testaf = testdf.withColumn("features",expr("strlisttoarray(features)"))

predict_df = model.transform(testaf)
predict_df.write.format("gor").save(f"{projectdir}/{resultdir}/testing_results.gord")

spark.stop()
spark._jvm.System.exit(0)
```

**Example 14:** Comparing good/bad labeling with VQSLOD and XGBoost

```
create #vqslod# = gor -p chr2 freezes/i2_37k_jg_v3/main/metadata/variant_qc_table.gorz | select 1-4,vqslod | calc posvqslod if(vqslod >= 2.41

create #giab# = gor -p chr2 user_data/hakon/freezes/test/i2_37k_giab_ff_v9_full_genome/v2/predictions_everything.gord
      | calc prob float(form(float(listtail(probability)),5,5))
| select 1-4,prob | calc posxgboost if(prob>=0.5,1,0);



create #paggr# = gor [#giab#] | select 1-4
| merge <(gor [#vqslod#] | select 1-4)
| varnorm -left #3 #4
| distinct
| calc snpindel if(len(ref)=len(alt),'snp','indel')
| varjoin -ic <(gor freezes/test/i2_37k_giab_ff_v9_full_genome/main/metadata/AF.gorz | where af >= 0.001)
| calc common if(overlapcount > 0,'comm','rare')

| varjoin -r  [#giab#]
| varjoin -r [#vqslod#]
| group chrom -gc posvqslod,posxgboost,snpindel,common -count;


create #all# = nor [#paggr#]
##commfilter##
| calc vqslod if(posvqslod=1,'good','bad')
| calc xgboost if(posxgboost=1,'good','bad')
| group -gc vqslod,xgboost -sum -ic allcount | rename sum_allcount allcount
| granno -sum -ic allcount
| calc f form(100.0*allcount/sum_allcount,4,2)
| hide sum_allcount;


create #snps# = nor [#paggr#]
##commfilter##
| where snpindel = 'snp'
| calc vqslod if(posvqslod=1,'good','bad')
| calc xgboost if(posxgboost=1,'good','bad')
| group -gc vqslod,xgboost -sum -ic allcount | rename sum_allcount allcount
| granno -sum -ic allcount
| calc f form(100.0*allcount/sum_allcount,4,2)
| hide sum_allcount;

create #indel# = nor [#paggr#]
##commfilter##
| where snpindel = 'indel'
| calc vqslod if(posvqslod=1,'good','bad')
| calc xgboost if(posxgboost=1,'good','bad')
| group -gc vqslod,xgboost -sum -ic allcount | rename sum_allcount allcount
| granno -sum -ic allcount
```

```
| calc f form(100.0*allcount/sum_allcount,4,2)
| hide sum_allcount;

def ##commfilter## = /* | where common = 'comm' */;
nor [#all#] | map -c vqslod,xgboost [#snps#] | map -c vqslod,xgboost [#indel#]
```

**Example 15:** Calculating the observed variant count as a function of samples number

```
create #segs# = gor -p chr2 freezes/test/i2_37k_giab_ff_v9_full_genome/main/metadata/AF.gorz | group 10000 -count | seghist 300000;

def #freeze# = freezes/test/i2_37k_giab_ff_v9_full_genome/main;
def #hgt# = #freeze#/variants.gord;
def #buckets# = #freeze#/buckets.tsv;

create #varminpn# = pgor -split <(gor [#segs#]) #hgt#
| csvsel -gc ref,alt -vs 1 -hide '0,3' #buckets# <(nor #buckets# | select pn) -tag PN
| group 1 -gc ref,alt -min -sc PN;

create #minnumcount# = nor [#varminpn#] | group -gc min_PN -count ;

create #temp# = SELECT allcount,
        SUM(allcount) over (ORDER BY -allcount ROWS BETWEEN unbounded preceding AND CURRENT ROW ) cumsum
 FROM  [#minnumcount#]
 ORDER  BY cumsum;

nor [#temp#]
```

**Example 16:** Performing concordance analysis between genotypes in two freezes, e.g. GOR fast-freeze and Sentieon-GATK

```
def #freeze1# = freezes/test/i2_37k_giab_ff_v9_full_genome/main;
def #freeze2# = freezes/test/i2_37k_giab_new_sention/main;
create #pns# = nor #freeze1#/buckets.tsv | select pn | inset -c pn #freeze2#/buckets.tsv /* | inset -c pn [#mypns#] */;

create #varstotest# = gor -p chr22 #freeze1#/metadata/AF.gorz | select 1-4;

create #segs# = gor [#varstotest#] | group 1000 -count | seghist 10000;

create #comp# = pgor -split <(gor [#segs#]) [#varstotest#] | varjoin -norm -ir #freeze1#/variants.gord
| csvsel -gc 3,4 -vs 1 -u 3 #freeze1#/buckets.tsv [#pns#]
| varjoin -norm -r <(gor #freeze2#/variants.gord | csvsel -gc ref,alt -vs 1 -u 3 #freeze2#/buckets.tsv [#pns#] )
| calc c listzip(fsvmap(values,1,'x',','),fsvmap(valuesx,1,'x',','))
| hide values,valuesx
| replace c listcount(c)
| split c
| colsplit c 3 x -s ';'
| rename x_1 gt1
| rename x_2 gt2
| rename x_3 count
| hide c
;

/* Seeing per variant concordance */
create #temp1# = gor [#comp#]
| where gt1 != 3 and gt2 !=3 and not(gt1=0 and gt2=0)
| group 1 -gc ref,alt,gt1,gt2 -sum -ic count
| calc x if(gt1!=gt2,sum_count,0)
| granno 1 -gc ref,alt -sum -ic sum_count,x
| calc f sum_count/sum_sum_count
| calc e sum_x/sum_sum_count
| hide x-sum_x
| sort 1 -c ref,alt,sum_count:n
| varjoin -r <(gor #freeze1#/metadata/AF.gorz | select 1-4,af)
| calc freqband if(af>=0.1,'>=0.1',if(af>=0.01,'>=0.01',if(af>=0.001,'>=0.001',if(af>=0.0001,'>=0.0001','lower'))))
| calc snpindel if(len(ref)=len(alt),'snp','indel')
| varjoin -r <(gor user_data/hakon/freezes/test/i2_37k_giab_ff_v9_full_genome/v2/predictions_everything.gord
   | calc Prob float(form(float(listtail(probability)),5,5)) | calc goodbad if(prob>=0.5,'good','bad') | select 1-4,goodbad)
| group 1 -gc ref,alt,e,af-;

create #temp2# = gor [#comp#]
| where gt1 != 3 and gt2 !=3
| group 1 -gc ref,alt,gt1,gt2 -sum -ic count
| calc x if(gt1!=gt2,sum_count,0)
| granno 1 -gc ref,alt -sum -ic sum_count,x
| calc f sum_count/sum_sum_count
| calc e sum_x/sum_sum_count
| hide x-sum_x
| sort 1 -c ref,alt,sum_count:n
| varjoin -r <(gor #freeze1#/metadata/AF.gorz | select 1-4,af)
| calc freqband if(af>=0.1,'>=0.1',if(af>=0.01,'>=0.01',if(af>=0.001,'>=0.001',if(af>=0.0001,'>=0.0001','lower'))))
```

```
| calc snpindel if(len(ref)=len(alt),'snp','indel')
| varjoin -r <(gor user_data/hakon/freezes/test/i2_37k_giab_ff_v9_full_genome/v2/predictions_everything.gord
    | calc Prob float(form(float(listtail(probability)),5,5)) | calc goodbad if(prob>=0.5,'good','bad') | select 1-4,goodbad)
| group 1 -gc ref,alt,e,af-;


create #base# = nor [#temp1#] | calc x round(if(e=0,-6,log(e))) | group -gc x -count
| map -c x <(nor [#temp2#] | calc x round(if(e=0,-6,log(e))) | group -gc x -count)
| map -c x <(nor [#temp1#] | where goodbad='good'| calc x round(if(e=0,-6,log(e))) | group -gc x -count
| map -c x <(nor [#temp2#] | where goodbad='good'| calc x round(if(e=0,-6,log(e))) | group -gc x -count));

nor [#base#] | granno -sum -ic allcount,allcountx,allcountxx,allcountxxx
| calc f allcount/sum_allcount
| calc fx allcountx/sum_allcountx
| calc fxx allcountxx/sum_allcountxx
| calc fxxx allcountxxx/sum_allcountxxx
| hide sum_*
| replace f* form(float(#rc)*100,6,2)+'%'
| calc band form(pow(10,x+0.5),1,7)+' - '+form(pow(10,x-0.5),1,7)
| select  x,band,allCount,f,allCountx,fx,allCountxx,fxx,allCountxxx,fxxx
```

### 2 Result Tables

Table 1: GIAB SNPs and InDels for chr22

| GIAB Sample | Num Vars | biallele TPR | GATK TPR | GOR TPR | biallele PPV | GATK PPV | GOR PPV | biallele F1 | GATK F1 | GOR F1 |
|---|---|---|---|---|---|---|---|---|---|---|
| HG001 | 47028 | 0.9906 | 0.9922 | 0.9921 | 0.9815 | 0.9824 | 0.9830 | 0.9860 | 0.9873 | 0.9875 |
| HG002 | 44382 | 0.9945 | 0.9949 | 0.9951 | 0.9506 | 0.9510 | 0.9514 | 0.9721 | 0.9724 | 0.9727 |
| HG003 | 45327 | 0.9942 | 0.9947 | 0.9947 | 0.9495 | 0.9490 | 0.9506 | 0.9713 | 0.9713 | 0.9722 |
| HG004 | 42167 | 0.9933 | 0.9938 | 0.9939 | 0.9441 | 0.9451 | 0.9456 | 0.9681 | 0.9688 | 0.9691 |
| HG006 | 44591 | 0.9896 | 0.9911 | 0.9914 | 0.9848 | 0.9853 | 0.9860 | 0.9872 | 0.9882 | 0.9887 |
| HG007 | 46646 | 0.9908 | 0.9912 | 0.9918 | 0.9830 | 0.9844 | 0.9844 | 0.9869 | 0.9878 | 0.9881 |
| NA12878 | 47040 | 0.9916 | 0.9930 | 0.9931 | 0.9819 | 0.9798 | 0.9823 | 0.9867 | 0.9864 | 0.9877 |

Table 2: GIAB SNPs for chr22

| GIAB Sample | Num Vars | biallele TPR | GATK TPR | GOR TPR | biallele PPV | GATK PPV | GOR PPV | biallele F1 | GATK F1 | GOR F1 |
|---|---|---|---|---|---|---|---|---|---|---|
| HG001 | 40671 | 0.9913 | 0.9935 | 0.9929 | 0.9807 | 0.9818 | 0.9821 | 0.9859 | 0.9876 | 0.9875 |
| HG002 | 39253 | 0.9953 | 0.9964 | 0.9959 | 0.9569 | 0.9575 | 0.9576 | 0.9757 | 0.9766 | 0.9764 |
| HG003 | 40212 | 0.9952 | 0.9960 | 0.9957 | 0.9572 | 0.9576 | 0.9584 | 0.9758 | 0.9764 | 0.9767 |
| HG004 | 37235 | 0.9940 | 0.9951 | 0.9946 | 0.9516 | 0.9533 | 0.9531 | 0.9723 | 0.9737 | 0.9734 |
| HG006 | 39217 | 0.9901 | 0.9926 | 0.9920 | 0.9842 | 0.9852 | 0.9856 | 0.9872 | 0.9889 | 0.9888 |
| HG007 | 40990 | 0.9914 | 0.9931 | 0.9925 | 0.9823 | 0.9841 | 0.9839 | 0.9868 | 0.9886 | 0.9882 |
| NA12878 | 40671 | 0.9916 | 0.9936 | 0.9932 | 0.9809 | 0.9795 | 0.9812 | 0.9862 | 0.9865 | 0.9872 |

Table 3: GIAB InDels for chr22

| GIAB Sample | Num Vars | biallele TPR | GATK TPR | GOR TPR | biallele PPV | GATK PPV | GOR PPV | biallele F1 | GATK F1 | GOR F1 |
|---|---|---|---|---|---|---|---|---|---|---|
| HG001 | 6357 | 0.9860 | 0.9842 | 0.9874 | 0.9872 | 0.9866 | 0.9886 | 0.9866 | 0.9854 | 0.9880 |
| HG002 | 5129 | 0.9885 | 0.9834 | 0.9887 | 0.9052 | 0.9032 | 0.9060 | 0.9450 | 0.9416 | 0.9455 |
| HG003 | 5115 | 0.9861 | 0.9841 | 0.9873 | 0.8926 | 0.8857 | 0.8932 | 0.9370 | 0.9323 | 0.9379 |
| HG004 | 4932 | 0.9884 | 0.9837 | 0.9888 | 0.8907 | 0.8871 | 0.8917 | 0.9370 | 0.9329 | 0.9378 |
| HG006 | 5374 | 0.9862 | 0.9808 | 0.9872 | 0.9886 | 0.9861 | 0.9892 | 0.9874 | 0.9834 | 0.9882 |
| HG007 | 5656 | 0.9869 | 0.9775 | 0.9867 | 0.9876 | 0.9862 | 0.9885 | 0.9873 | 0.9818 | 0.9876 |
| NA12878 | 6369 | 0.9918 | 0.9890 | 0.9923 | 0.9887 | 0.9820 | 0.9892 | 0.9903 | 0.9855 | 0.9907 |

Table 4: GIAB, good SNPs for chr22

| GIAB Sample | Num Vars | biallele TPR | GATK TPR | GOR TPR | biallele PPV | GATK PPV | GOR PPV | biallele F1 | GATK F1 | GOR F1 |
|---|---|---|---|---|---|---|---|---|---|---|
| HG001 | 39949 | 0.9989 | 0.9990 | 0.9990 | 0.9999 | 0.9998 | 0.9999 | 0.9994 | 0.9994 | 0.9994 |
| HG002 | 38774 | 0.9990 | 0.9991 | 0.9990 | 0.9841 | 0.9840 | 0.9841 | 0.9915 | 0.9915 | 0.9915 |
| HG003 | 39689 | 0.9992 | 0.9993 | 0.9992 | 0.9829 | 0.9827 | 0.9829 | 0.9910 | 0.9909 | 0.9910 |
| HG004 | 36692 | 0.9988 | 0.9990 | 0.9988 | 0.9820 | 0.9819 | 0.9820 | 0.9903 | 0.9904 | 0.9903 |
| HG006 | 38310 | 0.9989 | 0.9990 | 0.9991 | 1.0000 | 0.9999 | 1.0000 | 0.9995 | 0.9994 | 0.9995 |
| HG007 | 40149 | 0.9993 | 0.9994 | 0.9994 | 0.9999 | 0.9999 | 0.9999 | 0.9996 | 0.9997 | 0.9997 |
| NA12878 | 39949 | 0.9995 | 0.9996 | 0.9996 | 0.9999 | 0.9997 | 0.9999 | 0.9997 | 0.9997 | 0.9997 |

Table 5: GIAB, good InDels for chr22

| GIAB Sample | Num Vars | biallele TPR | GATK TPR | GOR TPR | biallele PPV | GATK PPV | GOR PPV | biallele F1 | GATK F1 | GOR F1 |
|---|---|---|---|---|---|---|---|---|---|---|
| HG001 | 5879 | 0.9952 | 0.9937 | 0.9957 | 0.9990 | 0.9990 | 0.9990 | 0.9971 | 0.9963 | 0.9974 |
| HG002 | 4727 | 0.9975 | 0.9917 | 0.9975 | 0.9576 | 0.9569 | 0.9576 | 0.9771 | 0.9740 | 0.9771 |
| HG003 | 4692 | 0.9966 | 0.9932 | 0.9968 | 0.9407 | 0.9404 | 0.9407 | 0.9678 | 0.9660 | 0.9679 |
| HG004 | 4551 | 0.9976 | 0.9921 | 0.9978 | 0.9441 | 0.9443 | 0.9441 | 0.9701 | 0.9676 | 0.9702 |
| HG006 | 4896 | 0.9955 | 0.9910 | 0.9959 | 0.9988 | 0.9986 | 0.9988 | 0.9971 | 0.9948 | 0.9973 |
| HG007 | 5148 | 0.9959 | 0.9877 | 0.9955 | 0.9982 | 0.9974 | 0.9982 | 0.9971 | 0.9926 | 0.9969 |
| NA12878 | 5886 | 0.9990 | 0.9973 | 0.9990 | 0.9997 | 0.9993 | 0.9997 | 0.9993 | 0.9983 | 0.9993 |

Table 6: All vars - whole genome

| GIAB Sample | Num Vars | GOR TPR | biallele TPR | GOR PPV | biallele PPV | GOR F1 | biallele F1 | GOR TP | biallele TP | GOR FN | biallele FN | GOR FP | biallele FP |
|---|---|---|---|---|---|---|---|---|---|---|---|---|---|
| HG001 | 3771313 | 0.9945 | 0.9935 | 0.9920 | 0.9911 | 0.9932 | 0.9923 | 374675 | 374674 | 208170 | 245660 | 302610 | 335100 |
| HG002 | 3722374 | 0.9950 | 0.9942 | 0.9674 | 0.9668 | 0.9810 | 0.9803 | 370059 | 370070 | 185100 | 216670 | 124869 | 127242 |
| HG003 | 3694081 | 0.9947 | 0.9939 | 0.9640 | 0.9634 | 0.9791 | 0.9784 | 367139 | 367143 | 194420 | 226480 | 137011 | 139549 |
| HG004 | 3719041 | 0.9949 | 0.9940 | 0.9647 | 0.9641 | 0.9796 | 0.9788 | 369678 | 369687 | 190100 | 221650 | 135184 | 137724 |
| HG006 | 3737516 | 0.9940 | 0.9930 | 0.9921 | 0.9915 | 0.9931 | 0.9922 | 371096 | 371124 | 222560 | 262760 | 294620 | 319990 |
| HG007 | 3754897 | 0.9937 | 0.9926 | 0.9921 | 0.9913 | 0.9929 | 0.9920 | 372696 | 372720 | 234710 | 276970 | 297060 | 326240 |
| NA12878 | 3772364 | 0.9951 | 0.9939 | 0.9918 | 0.9916 | 0.9934 | 0.9927 | 374940 | 374949 | 183920 | 228730 | 311570 | 318940 |

Table 7: All good vars - whole genome

| GIAB Sample | Num Vars | GOR TPR | biallele TPR | GOR PPV | biallele PPV | GOR F1 | biallele F1 | GOR TP | biallele TP | GOR FN | biallele FN | GOR FP | biallele FP |
|---|---|---|---|---|---|---|---|---|---|---|---|---|---|
| HG001 | 3701458 | 0.9986 | 0.9985 | 0.9997 | 0.9997 | 0.9992 | 0.9991 | 369608 | 369592 | 515600 | 553300 | 106000 | 106700 |
| HG002 | 3655963 | 0.9993 | 0.9993 | 0.9837 | 0.9837 | 0.9914 | 0.9914 | 365327 | 365324 | 258100 | 272100 | 606660 | 606680 |
| HG003 | 3628762 | 0.9991 | 0.9991 | 0.9813 | 0.9813 | 0.9901 | 0.9901 | 362539 | 362531 | 323200 | 344500 | 689580 | 689650 |
| HG004 | 3651998 | 0.9993 | 0.9992 | 0.9815 | 0.9815 | 0.9903 | 0.9903 | 364921 | 364918 | 266200 | 281700 | 687560 | 687610 |
| HG006 | 3658186 | 0.9991 | 0.9991 | 0.9998 | 0.9998 | 0.9995 | 0.9995 | 365481 | 365478 | 322700 | 340500 | 552000 | 559000 |
| HG007 | 3673624 | 0.9990 | 0.9990 | 0.9998 | 0.9998 | 0.9994 | 0.9994 | 366986 | 366984 | 358400 | 377500 | 610000 | 615000 |
| NA12878 | 3702345 | 0.9994 | 0.9994 | 0.9998 | 0.9998 | 0.9996 | 0.9996 | 370000 | 369996 | 225900 | 237600 | 736000 | 738000 |
